# Sexually dimorphic activation of innate antitumour immunity prevents adrenocortical carcinoma development

**DOI:** 10.1101/2022.04.29.489846

**Authors:** James J Wilmouth, Julie Olabe, Diana Garcia-Garcia, Cécily Lucas, Rachel Guiton, Florence Roucher-Boulez, Damien Dufour, Christelle Damon-Soubeyrand, Isabelle Sahut-Barnola, Jean-Christophe Pointud, Yoan Renaud, Adrien Levasseur, Igor Tauveron, Anne-Marie Lefrançois-Martinez, Antoine Martinez, Pierre Val

**Affiliations:** Institut GReD (Genetics, Reproduction and Development), CNRS UMR 6293, Inserm U1103, Université Clermont Auvergne, 28 Place Henri Dunant 63000 Clermont-Ferrand, France; Laboratoire de Biochimie et Biologie Moléculaire, UM Pathologies Endocriniennes, Groupement Hospitalier Est, Hospices Civils de Lyon, Bron, France; Univ Lyon, Université Claude Bernard Lyon 1, Lyon, France; Endocrinologie Diabétologie CHU Clermont Ferrand 58 rue Montalembert F63000 Clermont Fd France

**Keywords:** sexual dimorphism, cancer, macrophages, antitumour immunity, phagocytosis, senescence, senescence associated secretory phenotype, adrenocortical carcinoma, androgens, ZNRF3, WNT signalling

## Abstract

In contrast with most cancers, adrenocortical carcinomas (ACC) are more frequent in women than men, but the underlying mechanisms of this sexual dimorphism remain elusive. Homozygous deletion of the negative WNT pathway regulator *ZNRF3* is the most frequent alteration in ACC patients. Here, we show that Cre-mediated inactivation of *Znrf3* in steroidogenic cells of the mouse adrenal cortex is associated with sexually dimorphic tumour progression. Indeed, although most knockout female mice develop metastatic carcinomas over an 18 month-time course, adrenal hyperplasia gradually regresses in male knockout mice. This male-specific regression is associated with induction of senescence and recruitment of macrophages, which differentiate as active phagocytes that clear-out senescent preneoplastic cells. Macrophage recruitment is also observed in female mice. However, it is delayed and dampened compared to males, which allows for tumour progression. Interestingly, testosterone treatment of female knockouts is sufficient to induce senescence, recruitment of phagocytic macrophages and regression of hyperplasia. We further show that although macrophages are present within adrenal tumours at 18 months, MERTK^high^ active phagocytes are mostly found in indolent lesions in males but not in aggressive tumours in females. Consistent with our observations in mice, analysis of RNA sequencing data from the TCGA cohort of ACC shows that phagocytic macrophages are more prominent in men than women and associated with better prognosis. Altogether, these data establish that phagocytic macrophages prevent aggressive ACC development in male mice and suggest that they may play a key role in the unusual sexual dimorphism of ACC in patients.

## Introduction

Apart from reproductive tissues, cancer incidence and mortality are higher in males than females^1,2^. Adrenocortical carcinoma (ACC), which arises from steroidogenic cells of the adrenal cortex is one of the rare exceptions to this rule. Indeed, ACC female-to-male ratios range from 1.5 to 2.5:1 and women are generally diagnosed at a younger age (Fig S1A)^3–7^. Although the higher rate of steady state proliferation and more efficient adrenal cortex renewal in females ^5,8,9^ may play a role in sexually dimorphic tumorigenesis, the mechanisms underlying female prevalence of ACC remain elusive.

ACC is an aggressive cancer, with half of the patients presenting with metastatic disease at diagnosis. Overall, 5 year survival rates range between 16 and 47% and decrease to around 10% for metastatic patients^10^. In line with the steroidogenic activity of the adrenal cortex, ACC is associated with hormonal hypersecretion in more than 50% of patients^11^. A vast majority of secreting ACC produce excess glucocorticoids, but some tumours also produce sex steroids or a combination of both ^12^.

Radical surgical resection of ACC is the most effective therapeutic strategy for localized tumours, but the risk of recurrence remains high^12^. In patients with advanced inoperable or metastatic ACC, the adrenolytic compound mitotane, a derivate of the insecticide DDT, remains the standard of care, used as a single agent or in combination with an etoposide-doxorubicin-platin polychemotherapy^13–15^. Although these treatments can improve recurrence free survival, their benefit on overall survival is still debated^12,13,16–18^. Several phase I/II clinical trials of immune checkpoint inhibitors targeting PD1 and PD-L1, have also been conducted in ACC patients^19–22^. Unfortunately, these were associated with low response rates and have failed to improve patient outcome significantly. One potential reason for these modest results is the low level of lymphocyte infiltration in ACC^23^, which seems associated with local production of glucocorticoids^24^.

Understanding the molecular underpinnings of ACC pathogenesis is thus of utmost importance to develop novel therapeutic approaches. Large scale pan-genomic studies have identified homozygous deletion of *ZNRF3* as the most frequent genetic alteration in ACC^25,26^. This gene encodes a membrane E3 ubiquitin-ligase that inhibits WNT signalling by inducing ubiquitination and degradation of Frizzled receptors^27,28^. We previously showed that conditional ablation of *Znrf3* within steroidogenic cells of the adrenal cortex, resulted in moderate WNT pathway activation and adrenal zona fasciculata hyperplasia up to 6 weeks, suggesting that ZNRF3 was a potential tumour suppressor in the adrenal cortex^29^. However, we did not evaluate later stages of tumour progression.

Here, we show that tumour progression following ablation of *Znrf3* within steroidogenic cells of the adrenal cortex is sexually dimorphic. Whereas most female mice develop full-fledged metastatic carcinomas over an 18 month-time course, adrenal hyperplasia gradually regresses in male knockout mice. We show that male-specific regression of hyperplasia is associated with induction of senescence, recruitment of macrophages and differentiation of active phagocytes that clear out senescent steroidogenic cells. Although some degree of macrophage recruitment is observed in female mice, it is delayed and dampened compared to males, which allows for tumour progression. This phenomenon is dependent on androgens and can be triggered by testosterone treatment in females. Interestingly, even though macrophages are present within adrenal tumours at 18 months, active phagocytes, characterised by expression of the TAM receptor MERTK are mostly found in males but not females. Consistent with our observations in mice, analysis of RNA sequencing data from the TCGA cohort of ACC shows that phagocytic macrophages are more prominent in men than women and associated with better prognosis. Altogether, these data establish that phagocytic macrophages prevent aggressive ACC development in male mice and suggest that they may play a key role in the unusual sexual dimorphism of ACC in patients.

## Results

### Sexually dimorphic tumour progression in *Znrf3 cKO* adrenals

We previously showed that adrenal targeted ablation of *Znrf3* resulted in massive zona fasciculata hyperplasia at 6 weeks of age, but we did not evaluate the phenotype at later stages^29^. To gain further insight into the potential tumour suppressor function of ZNRF3 in the adrenal cortex, we conducted a kinetic analysis from 4 to 78 weeks (Fig 1). In female *Znrf3 cKO* mice (ZKO), adrenal weight increased progressively from 4 to 6 weeks and plateaued from 9 to 24 weeks. By 52 weeks, *Znrf3 cKO* average adrenal weight was still significantly increased and some adrenals were massively enlarged with weights up to 534mg. This trend was further amplified at 78 weeks with a majority of female *Znrf3 cKO* adrenals showing over a 10-fold increase in weight compared to controls (Fig 1A). This suggested malignant transformation of adrenals over time. Consistent with this idea, introduction of the mTmG reporter in the breeding scheme allowed identification of multiple micro and macro-metastases in the local lymph nodes, peritoneal cavity, liver and lungs of 75% of female *Znrf3 cKO* mice at 78 weeks (Fig 1B and Fig S1B). Histological analysis of adrenals that were associated with metastatic development (Fig 1C) showed complete disorganisation of the cortex that was mostly composed of densely packed small basophilic cells. This was associated with a significant increase in Ki67 labelling index (Fig 1C & D), although proliferation was rather heterogeneous throughout the tumour with areas showing up to 25% Ki67 labelling (Fig S1C). In contrast, in the few mutant mice where no metastases were found at 78 weeks (indolent ZKO), adrenals were largely hyperplastic, but cells retained a relatively normal morphology and Ki67 labelling was similar to control (Fig 1C). Altogether, these data suggested that ZNRF3 behaved as a classical tumour suppressor in female mice, its ablation resulting in a high frequency of aggressive adrenocortical carcinoma formation at 78 weeks. In sharp contrast, although male *Znrf3 cKO* adrenals were also larger at 4 and 6 weeks, adrenal weight steadily declined thereafter, almost returning to normal at 78 weeks (Fig 1E). This was associated with lack of metastatic progression (Fig 1F), benign histology and low Ki67 labelling index (Fig 1G & H), although some patches of higher proliferation could be detected in some adrenals (Fig S1C). This suggested that overall tumour development was rapidly blunted in males, although the initial hyperplastic phase was equivalent to females.

**Figure 1.**
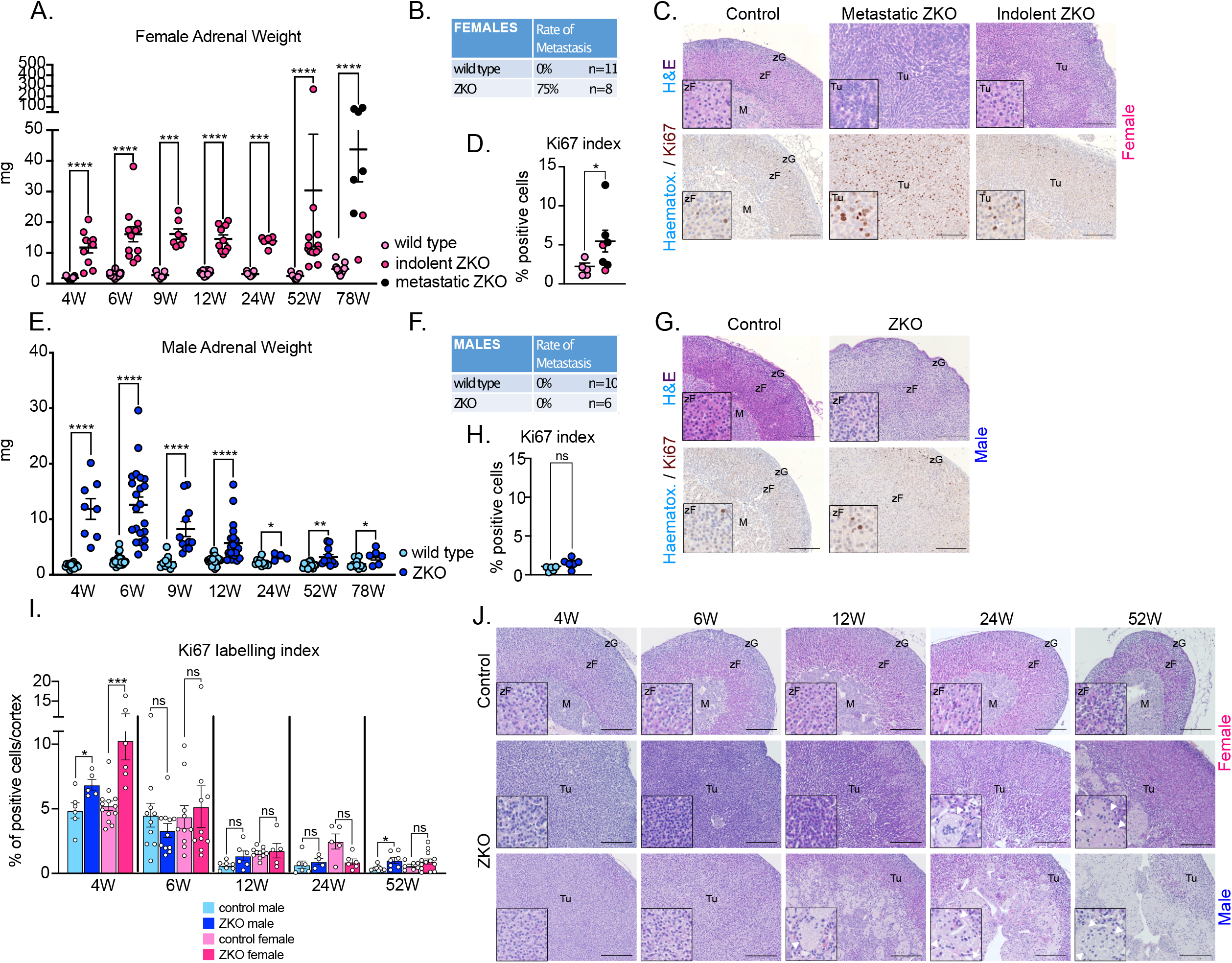
Sexually dimorphic tumour progression in *Znrf3 cKO* adrenals. **A-** Female adrenal weights measured from 4 to 78 weeks in wild-type and *Znrf3 cKO* (ZKO) adrenals. **B-** Rate of metastasis in 78-week-old *Znrf3 cKO* females. **C-** Histology (upper panels) and immunohistochemical analysis of Ki67 expression (lower panels) in 78-week-old female controls, *Znrf3 cKO* adrenals associated with metastasis formation or indolent *Znrf3 cKO* adrenals. **D-** Quantification of the Ki67 proliferation index as the ratio of positive cells over total nuclei in the cortex of 78-week-old control and *Znrf3 cKO* females. **E-** Male adrenal weights measured from 4 to 78 weeks in wild-type and *Znrf3 cKO* (ZKO) adrenals. **F-** Rate of metastasis in 78-week-old *Znrf3 cKO* males. **G-** Histology (upper panels) and immunohistochemical analysis of Ki67 expression (lower panels) in 78-week-old male controls and *Znrf3 cKO* adrenals. **H-** Quantification of the Ki67 proliferation index as the ratio of positive cells over total nuclei in the cortex of 78-week-old control and Znrf3 cKO males. **I-** Kinetic analysis of the Ki67 proliferation index from 4 to 52 weeks in male and female control and *Znrf3 cKO* adrenals. **J-** Kinetic analysis of the histological phenotype from 4 to 52 weeks in male and female control and *Znrf3 cKO* adrenals. Arrowheads in insets show multinucleated giant cells that accumulate in the inner cortex of mutant male mice and to a lesser extent mutant female mice. M: medulla; zF: zona fasciculata; zG zona glomerulosa; Tu: tumour. Scale bar = 200μm. Graphs represent mean +/- SEM. Statistical analyses in A, D, E, H and I were conducted by Mann-Whitney tests. ns : not significant; * p<0.05; ** p<0.01; *** p<0.001; **** p<0.0001.

To further gain insight into this sexually dimorphic phenotype, we evaluated proliferation from 4 to 52 weeks. Analysis of Ki67 labelling index showed that following an early significant increase, both males and females had a rapid arrest in proliferation at 6 weeks (Fig 1I & Fig S1D). This remained at low levels up to 52 weeks, although male adrenals displayed a mild but significant rebound at this stage (Fig 1I). The steady decline in adrenal weight, although proliferation in male knockout adrenals was comparable to controls after 4 weeks, suggested that an active mechanism counteracted tumour progression in males. Surprisingly though, there was no increase in apoptosis, measured by cleaved caspase 3 staining, in either female or male adrenals at 6 and 12 weeks (Fig S1E). To try to further understand the sexually dimorphic phenotype, we conducted a careful kinetic evaluation of adrenal histology. This showed a similar hyperplastic phenotype in males and females at 4 and 6 weeks (Fig 1J). Hyperplasia progressed in females with accumulation of small basophilic cells that composed most of the gland by 52 weeks (Fig 1J). Strikingly, starting at 12 weeks, we observed progressive thinning of the cortex (eosinophilic cells) and concomitant appearance and expansion of multinucleated giant cells (MGCs containing up to 12 nuclei per cell) that progressively took over a large proportion of the male *Znrf3 cKO* gland (up to 40%) (Fig 1J). In females some MGCs were also observed. However, they were first visible at 24 weeks and only represented a small proportion of the gland, even at 52 weeks (Fig 1J). Interestingly, MGCs were reminiscent of fused macrophages that are observed in granulomatous inflammatory diseases, which suggested a potential involvement of innate immune cells in preventing tumour progression in male *Znrf3 cKO* adrenals.

### Regression in male *Znrf3 cKO* adrenals is correlated with macrophage infiltration and fusion

To further gain insight into the underpinnings of the regression phenomenon, we analysed global gene expression by bulk RNA sequencing of control and *Znrf3 cKO* male adrenals at 4, 6 and 12 weeks. Gene set enrichment analysis (GSEA) of the RNA sequencing data using the C5 Gene Ontology database (MSigDB) showed that at 12 weeks, the 34 most significantly enriched gene sets were all related with immune response and inflammation (Fig 2A). Most of these gene sets were either not (FDR >0.05) or negatively enriched at 4 weeks and showed an intermediate enrichment score at 6 weeks. This suggested that ablation of Znrf3 resulted in the progressive establishment of a proinflammatory environment. Consistent with this idea, a large number of proinflammatory cytokines and chemokines genes were progressively upregulated at 6 and 12 weeks (Fig 2B and Fig S2A). Establishment of an inflammatory environment was further evaluated by immunohistochemistry for the pan leukocyte marker CD45. In control male adrenals, a few CD45-positive cells were found scattered throughout the cortex. Four-week-old male *Znrf3 cKO* adrenals were quite similar to controls, although more mononucleated leukocytes were present in the inner cortex. At 6 and 12 weeks, the number of CD45 positive cells dramatically increased in KO adrenals (Fig 2C and Fig S2B). These comprised both mononuclear cells (stars) and the multinucleated giant cells (arrowheads) that accumulated in the inner cortex (Fig 2C). To further identify the immune cell types that composed the infiltrate, we deconvoluted RNA sequencing data with CibersortX using immune cell signatures from ImmuCC (Fig 2D) and mMCP (Fig S2C). Both approaches showed a significant increase in macrophages populations, which represented 63% of all immune populations at 12 weeks. This was further confirmed by GSEA, showing a highly significant positive enrichment of multiple macrophages signatures at 12 weeks (Fig 2E) and by RTqPCR showing a progressive accumulation of the macrophage marker transcripts *Cd68, Adgre1* and *Cd11b* (Fig S2D). Altogether, these data strongly suggested that regression of adrenal cortex hyperplasia in *Znrf3 cKO* males was associated with establishment of a proinflammatory environment and massive recruitment of macrophages.

**Figure 2.**
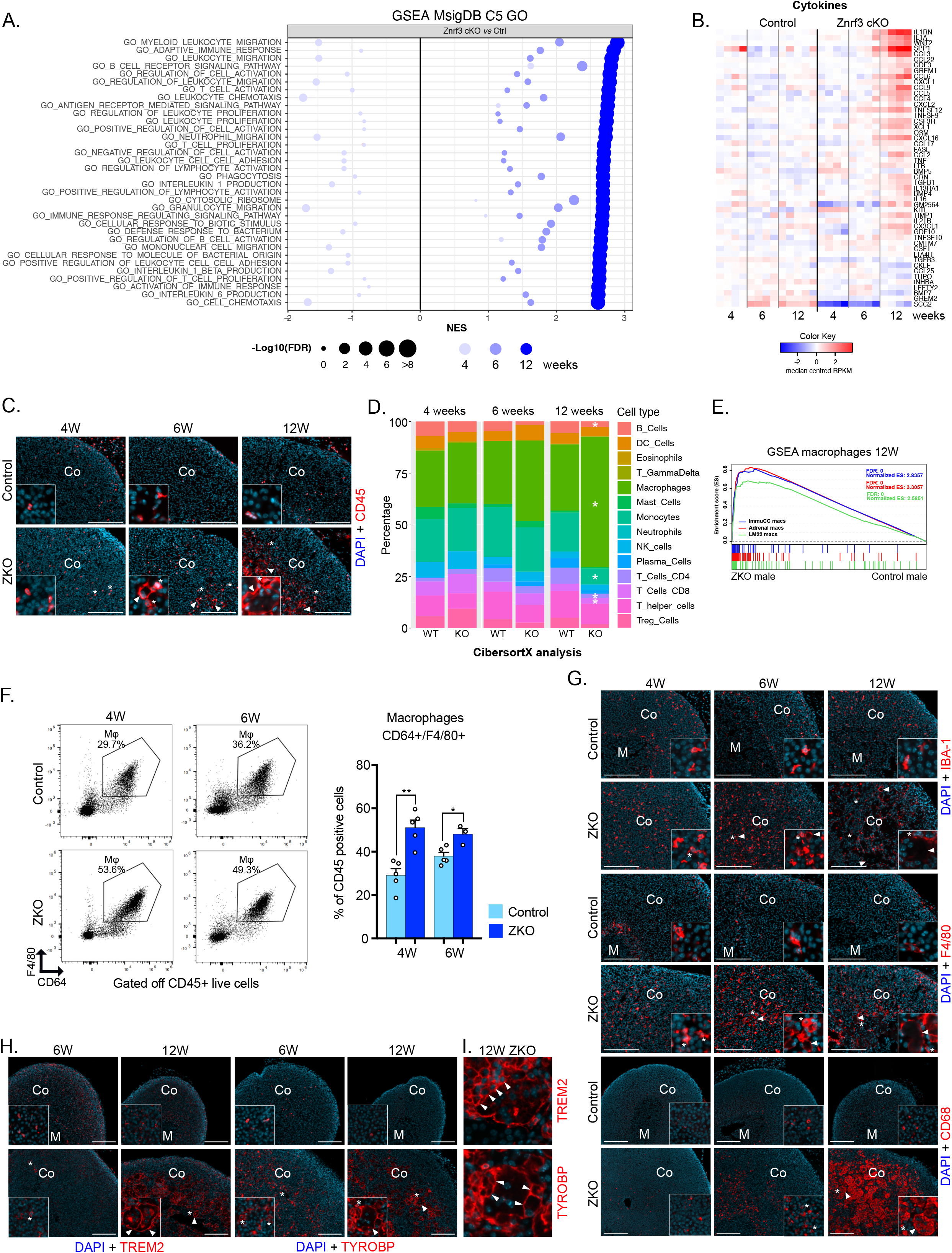
Regression in male *Znrf3 cKO* adrenals is correlated with macrophages infiltration and fusion. **A-** Gene Set Enrichment Analysis (GSEA) of gene expression data (RNA sequencing) from 4, 6 and 12-week-old control and *Znrf3 cKO* male adrenals. The plot represents the 35 gene sets from the C5 Gene Ontology database (MSigDB), with the highest enrichment score in *Znrf3 cKO* adrenals compared with controls at 12 weeks. **B-** Heatmap representing median-centred expression of cytokines/chemokines-coding genes in control and *Znrf3 cKO* adrenals at 4, 6 and 12 weeks. Only genes significantly deregulated at 12 weeks (FDR<0.05) are represented. They are sorted by decreasing Log2-fold-change. **C-** Immunohistochemical analysis of CD45 expression in adrenals from control and *Znrf3 cKO* (ZKO) mice at 4, 6 and 12 weeks. Arrowheads show multinucleated giant cells. Stars show mononucleated leukocytes. **D-** Stacked bar plots representing immune cell populations deconvoluted using CibersortX and the LM22 expression matrix, from gene expression data in control and *Znrf3 cKO* adrenals at 4, 6 and 12 weeks. **E-** Enrichment analysis (GSEA) of macrophages signatures derived from ImmuCC, LM22, and single cell RNA sequencing of mouse adrenals, in 12-week-old male *Znrf3 cKO* adrenals compared to controls. **F-** Left, representative dot plots of flow cytometry analysis of macrophages infiltration in 4 and 6-week-old control (top panels) and *Znrf3 cKO* (bottom panels) adrenals. Macrophages were defined as CD45^+^/CD64^+^/F4/80^+^ live cells. Right panel, quantification of flow cytometry experiments. **G-** Immunohistochemical analysis of pan-macrophages markers IBA-1, F4/80 and CD68 in 4, 6 and 12-week-old control and *Znrf3 cKO* adrenals. Arrowheads show multinucleated giant cells. Stars show mononucleated macrophages. **H-** Immunohistochemical analysis of macrophage fusion-associated markers TREM2 and TYROBP in 6 and 12-week-old control and *Znrf3 cKO* adrenals. Arrowheads show multinucleated macrophages. Stars show mononucleated macrophages. **I-** High-magnification images of TREM2 and TYROBP staining showing fusion of mononucleated with multinucleated macrophages in *Znrf3 cKO* adrenals at 12 weeks (arrowheads). Co: cortex; M: medulla; scale bar =200μm. Graphs represent mean +/- SEM. Statistical analyses in F, were conducted by Mann-Whitney tests. * p<0.05; ** p<0.01.

To further confirm the nature of infiltrating cells, adrenals from control and *Znrf3 cKO* males were dissociated and analysed by flow cytometry (Fig 2F and Fig S2E). In wild-type adrenals, CD64^+^/F4/80^+^ macrophages represented ~30 to 36% of all live CD45^+^ cells at 4 and 6 weeks. In *Znrf3 cKO* adrenals, this proportion was significantly increased up to ~49%-54% at these two stages, demonstrating increased macrophage infiltration as early as 4 weeks. Flow cytometry analyses further showed that at 4 weeks, almost 80%of CD45^+^/CD64^+^ macrophages co-expressed the M1 markers CD38 and MHC-II, together with the M2 marker CD206, both in wild-type and *Znrf3 cKO* adrenals (Fig S3A). Although there was a very mild but significant increase in both MHC-II^+^/CD206^+^ and CD38^+^/CD206^+^ double-positive macrophages in 6-week *Znrf3 cKO* adrenals, there was no significant difference in either M1 or M2 macrophages proportions, following ablation of *Znrf3* at the two analysed stages (Fig S3A). RTqPCR (Fig S3B) and RNA sequencing analyses (Fig S3C-D) further confirmed deregulation of both M1 and M2 markers in *Znrf3cKO* adrenals, indicating that infiltrating macrophages had mixed M1 and M2 characteristics at 4 and 6 weeks.

Unfortunately, the majority of CD45^+^ MGCs that accumulated from 12 weeks onward, had a cell diameter larger than 40 μm, which precluded their characterisation by flow cytometry (Fig S4A). To further characterise immune infiltration during the regression period, we thus resorted to immunohistochemical analysis. Staining with pan-macrophages markers IBA-1 and F4/80 confirmed progressive infiltration from 4 to 12 weeks (Fig 2G). Interestingly, although mononuclear cells appeared equivalently labelled by both IBA-1 and F4/80, IBA-1 staining of MGCs was weak compared to F4/80 (Fig 2G). However, MGCs displayed high levels of cytoplasmic CD68 staining, suggesting that they were derived from the fusion of mononuclear macrophages (Fig 2G). Macrophage fusion has been shown to rely on TREM2, an activating receptor of the Ig-superfamily and on TYROBP/DAP12, its transmembrane signalling adaptor 30,31. Interestingly, expression of *Trem2* and *Tyrobp/Dap12* was strongly increased in RTqPCR at 12 weeks (Fig S4B) and IHC analyses showed a strong up-regulation of both TREM2 and TYROBP protein accumulation in MGCs (Fig 2H). High magnification images further showed TREM2/TYROBP-positive mononuclear macrophages actively fusing with MGCs (Fig 2I, arrowheads).

Altogether, this suggested that *Znrf3* ablation in steroidogenic cells resulted in macrophage infiltration and fusion to form MGCs in male adrenals.

### Infiltrating macrophages actively phagocytose steroidogenic cells

Macrophages have been suggested to play a role in the early response to oncogenic insult, by clearing out preneoplastic cells^7,32^. Interestingly, GSEA of RNA sequencing data showed a progressive significant enrichment of gene sets associated with phagocytosis and clearance of apoptotic cells in male *Znrf3 cKO* adrenals, suggesting a potential role of phagocytosis in regression of hyperplasia (Fig 3A). Phagocytosis involves chemotaxis of macrophages towards target cells that express “find-me” signals and recognition of target cells through “eat-me” signals that can be received directly by phagocytic receptors, or indirectly after opsonization. Detailed analysis of RNA sequencing data showed significant up-regulation of genes coding the potential “find-me”chemokine CX3CL1^33^, and of the GPR132/G2A and P2RY2/P2RY6 metabotropic receptors that recognize lysophosphatidylcholine (GPR132)^34^ and nucleotides (P2RY2/P2RY6)^35^, released by target cells (Fig 3B). Among potential “eat-me” signals, we found significant overexpression of C1Q complement components *C1qa, C1qb* and *C1qc* which have been shown to decorate the surface of apoptotic cells to target them for phagocytosis ^35,36^ (Fig 3B), and of *Slamf7*, which is involved in phagocytosis of hematopoietic tumour cells ^37^. There was also upregulation of the gene coding MFGE8, which opsonizes apoptotic cells and is recognised by the integrin receptors α_v_β_3_ and α_v_β_5_ at the membrane of macrophages ^38,39^ (Fig 3B & 3C). Interestingly, TREM2 and TYROBP, which we found overexpressed both at the mRNA and protein level (Fig 2G, Fig 3B & Fig S4B), can also be involved in the phagocytic process through recognition of lipids and ApoE-opsonized cells ^38,40,41^. Among the three TAM receptor tyrosine kinases, which play a central role in phagocytosis (TYRO3, MERTK, AXL)^35,42^, *Mertk* was expressed at high levels and showed the most significant upregulation in *Znrf3 cKO* adrenals (Fig 3B &C). Although there was no upregulation of *Gas6* and *Pros1*, the natural TAM receptors ligands ^35^, there was a strong overexpression of *Lgals3* (27-fold), which encodes Galectin-3, a phosphatidylserine-independent MERTK-specific opsonin ^38,43^ (Fig 3B). This was further confirmed by RTqPCR (Fig 3C), suggesting that engagement of MERTK by Galectin-3 may trigger phagocytosis of *Znrf3 cKO* hyperplastic steroidogenic cells.

**Figure 3.**
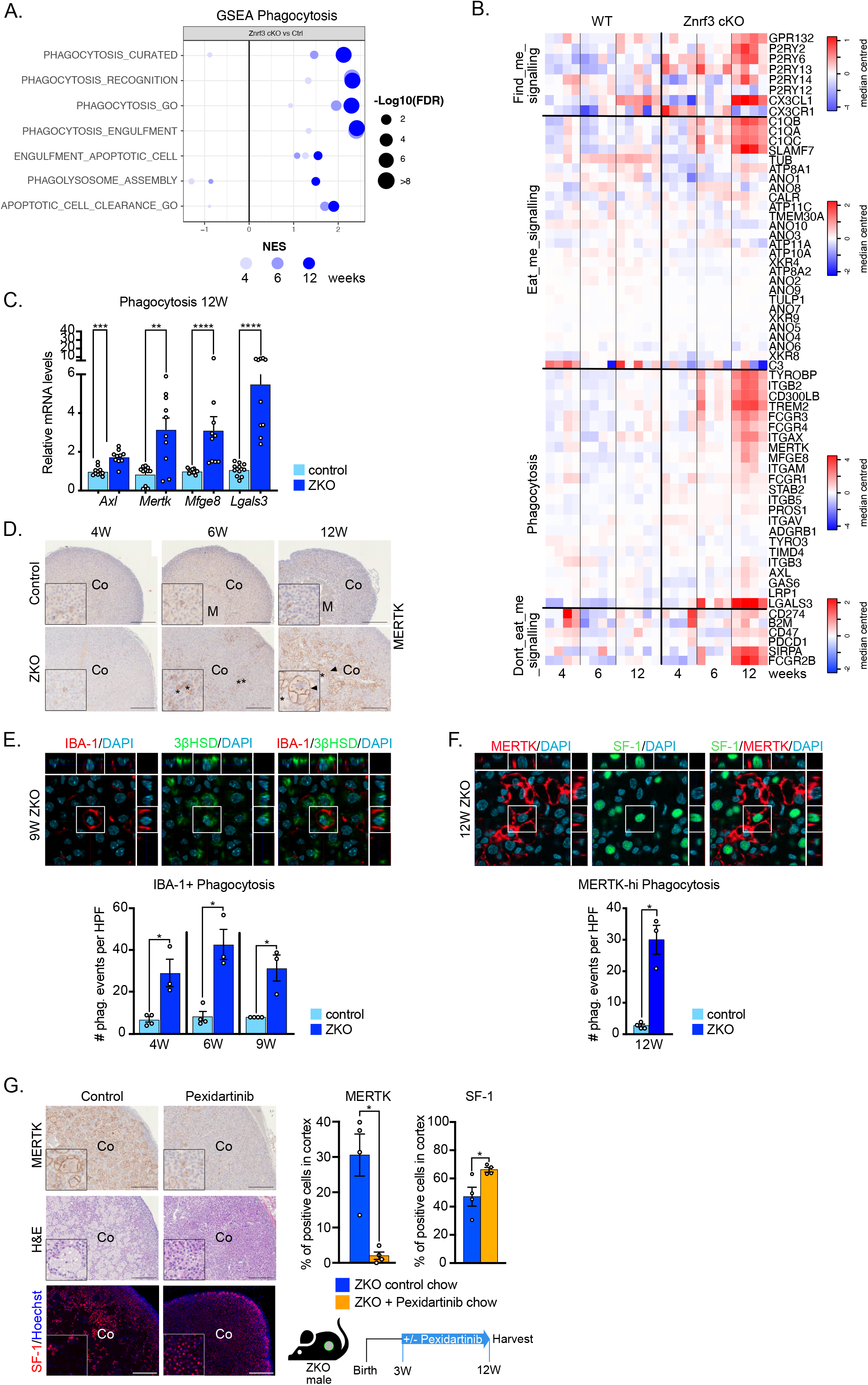
Infiltrating macrophages actively phagocytose steroidogenic cells. **A-** Gene Set Enrichment Analysis (GSEA) of gene expression data (RNA sequencing) from 4, 6 and 12-week-old control and *Znrf3 cKO* male adrenals. The plot represents enrichment of phagocytosis/efferocytosis gene sets in *Znrf3 cKO* male adrenals compared with controls. **B-** Heatmap representing median-centred expression of key regulators of the phagocytic pathway in control and *Znrf3 cKO* male adrenals at 4, 6 and 12 weeks. **C-** RTqPCR analysis of the expression of phagocytosis-associated genes in control and *Znrf3 cKO* male adrenals at 12 weeks. **D-** Immunohistochemical analysis of the expression of the phagocytosis receptor MERTK in control and *Znrf3 cKO* male adrenals at 4, 6 and 12 weeks. Arrowheads show multinucleated macrophages. Stars show mononucleated macrophages. **E-F** Evaluation of phagocytosis by immunohistochemistry for 3bHSD (steroidogenic cells) and IBA-1 (E) or SF-1 (steroidogenic cells) and MERTK (F). Images were acquired by confocal microscopy and phagocytic events were counted when steroidogenic markers were found within the boundaries of macrophages markers along the Z-stack. Panels show representative zoomed-in images (x120) in 9-week-old (IBA-1) and 12-week-old (MERTK) *Znrf3 cKO* adrenals. White boxes show phagocytic events on the 2D projection of Z-stack and within the orthogonal projections (side images). Bottom graphs represent quantification of phagocytic events on 10 high power fields (HPF, x40) per individual mouse from 4 to 9 weeks (IBA-1^+^ phagocytosis) and at 12 weeks (MERTK-hi phagocytosis). **G-** Immunohistochemical (MERTK & SF-1) and histological (H&E) analysis of *Znrf3 cKO* male mice that received control or Pexidartinib-enriched chow from 9 to 12 weeks (left panels). Percentages of MERTK-positive and SF-1-positive cells, relative to total cortical cell numbers (DAPI^+^) are displayed on the graphs (right panel). Co: cortex; M: medulla; scale bar =200μm. Graphs represent mean +/- SEM. Statistical analyses in C, E, F and G were conducted by Mann-Whitney tests. * p<0.05; ** p<0.01; *** p<0.001; **** p<0.0001.

To further gain insight into a potential phagocytic process in *Znrf3 cKO* adrenals, we analysed expression of the TAM receptor MERTK by IHC. Although some positive cells were found in wild-type adrenals, they were rather scarce and expressed low levels of MERTK (Fig 3D). In contrast, increased numbers of mononuclear MERTK^high^ cells were found in *Znrf3 cKO* adrenals as early as 6 weeks (Fig 3D). Most of these cells also stained for IBA-1, confirming their macrophage identity (Fig S4C). At 6 and 12 weeks, the number of mononuclear MERTK-high cells dramatically increased in *Znrf3 cKO* (Fig3D & Fig S4C). Interestingly, multinucleated fused macrophages expressed very high levels of MERTK (Fig 3D), which was associated with reduced IBA-1 expression (Fig S4C). MERTK^high^ and in particular, fused macrophages, were also positive for TREM2 (Fig S4D). However, TREM2 was expressed in a larger number of macrophages, including mononucleated MERTK^-^ macrophages (Fig S4D). Altogether, this suggested that macrophages infiltration in *Znrf3 cKO* male adrenals was associated with differentiation into active phagocytes.

To test this hypothesis, we evaluated phagocytosis by confocal microscopy. For this, we colocalised expression of 3βHSD and SF-1, two markers of steroidogenic cells with IBA-1 (from 4 to 9 weeks) and MERTK (at 12 weeks). We then counted 3βHSD and SF-1 positive cells that were found within the boundaries of IBA-1^+^ or MERTK^high^ macrophages throughout the confocal Z-stack (Fig 3E). A few IBA-1^+^ macrophages contained 3βHSD positive cells in control adrenals at 4, 6 and 9 weeks, indicating that phagocytosis of steroidogenic cells was taking place at homeostasis in the adrenal (Fig 3E). Interestingly, the number of phagocytic IBA-1^+^ cells was markedly increased in *Znrf3 cKO* adrenals at these three timepoints (Fig 3E), indicating that mononuclear IBA-1^+^ macrophages were actively involved in phagocytosis of *Znrf3 cKO* steroidogenic cells. Increased phagocytosis was also observed for MERTK^high^ macrophages at 12 weeks (Fig 3F).

Altogether, these data show that both IBA-1^+^ and MERTK^high^ macrophages are involved in a dramatic increase in phagocytosis of mutant steroidogenic cells in male *Znrf3 cKO* adrenals.

To further confirm the key role of macrophages in regression of adrenal hyperplasia, we depleted macrophages using a diet enriched with 290mg/kg Pexidartinib, a pharmacological inhibitor of CSF1R. This tyrosine kinase receptor plays a central role for survival of macrophages within their tissue niches, through stimulation by CSF1 and/or IL-34. Consistent with the key function of CSF1R, flow cytometry analyses showed that 1 week of pexidartinib chow was sufficient to deplete almost all CD45^+^/CD64^+^/F4/80^+^ macrophages within the adrenal cortex of control male mice (Fig S4E). We then evaluated the impact of macrophages depletion in male *Znrf3 cKO* mice by feeding them with standard chow or Pexidartinib chow from 3 to 12 weeks (Fig 3G). This resulted in a very strong decrease in the number of IBA-1^+^ (Fig S4F) and MERTK^high^ macrophages (Fig 3G) in IHC analyses. Consistent with these findings, H&E staining showed a remarkable decrease in the number of fused macrophages and concomitant expansion of presumptive eosinophilic steroidogenic cells (Fig 3G). This was further confirmed by a significant increase in SF-1 positive cells in the cortices of pexidartinib-treated mice (Fig 3G) and an inverse correlation between MERTK-positive and SF-1-positive cells (Fig S4G).

Altogether, these data show that *Znrf3* ablation induces sustained recruitment of IBA1^+^ and MERTK^high^ macrophages, which results in phagocytic clearance of mutant steroidogenic cells and regression of adrenal hyperplasia in male mice.

### Recruitment of phagocytic macrophages is delayed in females

In contrast with males, female *Znrf3 cKO* adrenals progress from hyperplasia at 4 weeks to development of full-fledged metastatic carcinomas at 78 weeks (Fig 1). Interestingly, analysis of the overall mononuclear macrophage population by IHC for IBA-1, showed increased recruitment of IBA-1-positive macrophages in *Znrf3 cKO* females from 4 to 52 weeks (Fig 4A). However, counting of IBA-1-positive cells suggested that macrophages recruitment was milder than in males from 4 to 12 weeks (Fig 4A&B). This was confirmed by GSEA (Fig 4C), showing a robust enrichment in macrophages signatures in male knockouts compared with female knockouts at 12 weeks and RTqPCR analyses of *Cd68, Adgre1* and *Cd11b* (Fig S5A). Milder inflammatory response in female knockouts was also confirmed by the absence of cytokine signature enrichment at 12 weeks, compared with male knockouts (Fig 4D). By 24 and up to 52 weeks, the number of IBA-1-positive cells significantly increased in female *Znrf3 cKO* adrenals, which was accompanied by a mild but significant increase in mRNA accumulation of *Adgre1* at 24 weeks and *Cd68* at 52 weeks (Fig 4A-B &Fig S5A). However, this was still not associated with enrichment of cytokines (Fig S5B). Altogether, this showed that macrophage recruitment was delayed in female *Znrf3 cKO* adrenals and was not associated with robust inflammation. In males, regression of hyperplasia is associated with fusion of mononuclear macrophages to form MGCs (Fig 3). Whereas fused macrophages were already present in large numbers in 12 weeks *Znrf3 cKO* males, they did not appear before 24 weeks in females (Fig 4E). Consistent with delayed fusion, fused macrophages harboured less nuclei (Fig S5C) and were smaller than in males at this stage (Fig S5D). In male *Znrf3 cKO* adrenals, acquisition of high phagocytic capacities is associated with infiltration of MERTK^high^ macrophages as early as 6 weeks (Fig 3D & Fig 4G). In contrast, these were scarce until 24 weeks in female *Znrf3 cKO* adrenals (Fig 4F-G). They mostly represented fused macrophages (Fig 4F) and were only significantly increased in numbers at 52 weeks (Fig 4G). This suggested that phagocytosis of hyperplastic mutant cells may be impaired in female knockouts. Indeed, although there was trend for increased phagocytosis by IBA-1^+^ macrophages, it did not reach significance, from 4 to 9 weeks (Fig 4H). Furthermore, the rate of phagocytosis was much lower than in males, barely reaching 12 events per high power field in female knockouts, compared with over 40 in male knockouts (Fig 3E). The low phagocytic capacity in females was even more evident when analysed within MERTK^high^ macrophages at 12 weeks (Fig 4H). This was supported by the lack of enrichment of phagocytosis-related gene signatures at any time point (Fig 4I), which was further confirmed by RTqPCR at 12 weeks (Fig 4J).

**Figure 4.**
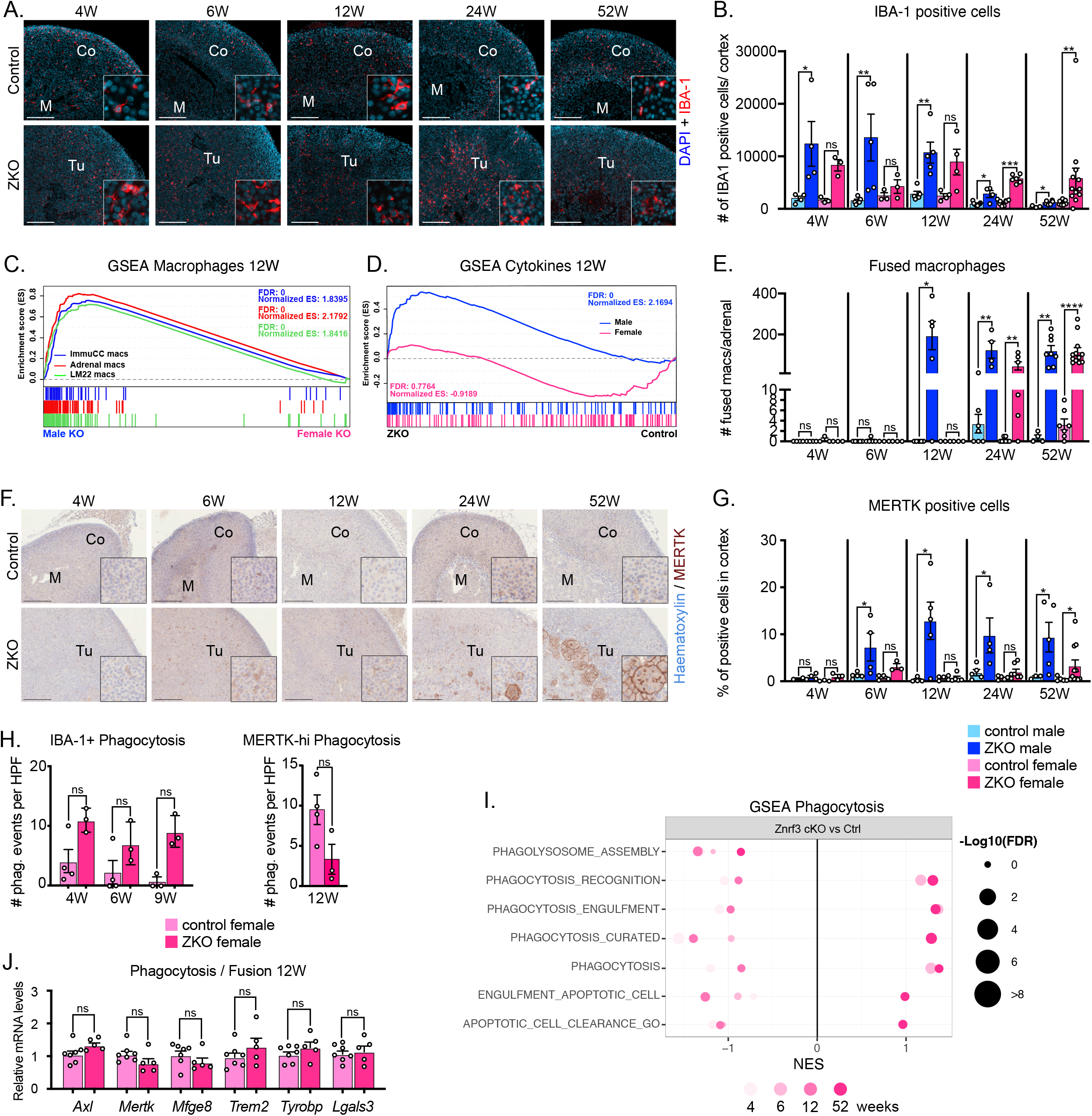
Recruitment of phagocytic macrophages is delayed in females. **A-** Immunohistochemical analysis of IBA-1 expression in female control and *Znrf3 cKO* adrenals from 4 to 52 weeks. **B-** Quantification of the IBA-1 index as the ratio of IBA-1-positive cells over total nuclei in the cortex of control and *Znrf3 cKO* male (blue) and female (pink) mice from 4 to 52 weeks. **C-** GSEA of macrophages gene sets in male *Znrf3 cKO* compared with female *Znrf3 cKO* adrenals at 12 weeks. **D-** GSEA of the cytokine gene set in *Znrf3 cKO* males and females compared with their respective control adrenals at 12 weeks. **E-** Quantification of the number of fused macrophages (at least 2 nuclei) in control and *Znrf3 cKO* male and female adrenals from 4 to 52 weeks. **F-** Immunohistochemical analysis of MERTK expression in control and *Znrf3 cKO* female adrenals from 4 to 52 weeks. **G-** Quantification of the MERTK^+^ index as the ratio of MERTK-positive cells over total nuclei in the cortex of control and *Znrf3 cKO* male (blue) and female (pink) mice from 4 to 52 weeks. **H-** Quantification of phagocytic events following immunohistochemistry for IBA-1 and 3bHSD (IBA-1^+^ phagocytosis) or MERTK and SF-1 (MERTK^high^ phagocytosis) in control and *Znrf3 cKO* females from 4 to 12 weeks. Quantification was performed on 10 high power fields (HPF, x40) per individual mouse. **I-** Gene Set Enrichment Analysis (GSEA) of gene expression data (RNA sequencing) from 4, 6, 12 and 52-week-old control and *Znrf3 cKO* female adrenals. The plot represents enrichment of phagocytosis/efferocytosis gene sets in *Znrf3 cKO* female adrenals compared with controls. **J-** RTqPCR analysis of the expression of phagocytosis and macrophages fusion-associated genes in control and *Znrf3 cKO* female adrenals at 12 weeks. Co: cortex; Tu: tumor; Scale bar = 200μm. Graphs represent mean +/- SEM. Statistical analyses in B, E, G, H and J were conducted by Mann-Whitney tests. ns: not significant; * p<0.05; ** p<0.01; *** p<0.001; **** p<0.0001.

Altogether, these data strongly suggest that delayed recruitment and impaired function of phagocytic macrophages allows progression of hyperplasia in *Znrf3 cKO* females.

### Androgens are sufficient to trigger early recruitment of phagocytic macrophages and regression of hyperplasia

Sexually dimorphic phenotypic differences in phagocytic macrophage recruitment and regression of hyperplasia occur between 6 and 12 weeks, which coincides with onset of puberty in mice. To evaluate a potential contribution of androgens to this phenomenon, *Znrf3 cKO* females were implanted with placebo or testosterone pellets from 4 to 12 weeks and their adrenals were then harvested (Fig 5A). As expected, placebo treated female adrenals were almost completely devoid of MERTK^high^ macrophages (Fig 5B-C). In sharp contrast, testosterone-treated females displayed massive infiltration of both mononuclear and fused MERTK^high^ macrophages, which was almost equivalent to 12-week-old males (Fig 5B-C). Infiltration of macrophages was further confirmed by RTqPCR showing increased expression of *Cd68, Adgre1* and *Cd11b*, following androgen treatment (Fig 5D). Interestingly, RTqPCR analysis of phagocytosis-associated gene expression also showed increased accumulation of *Axl, Mertk, Mfge8, Trem2, Tyrobp* and *Lgals3*, suggesting that testosterone treatment stimulated recruitment of phagocytic macrophages (Fig 5E). Consistent with this hypothesis, testosterone treatment was associated with a marked decrease in *Znrf3 cKO* female adrenal weight, which returned to control levels (Fig 5F). Altogether, these experiments show that androgens are sufficient to induce recruitment of phagocytic macrophages, which results in regression of hyperplasia.

**Figure 5.**
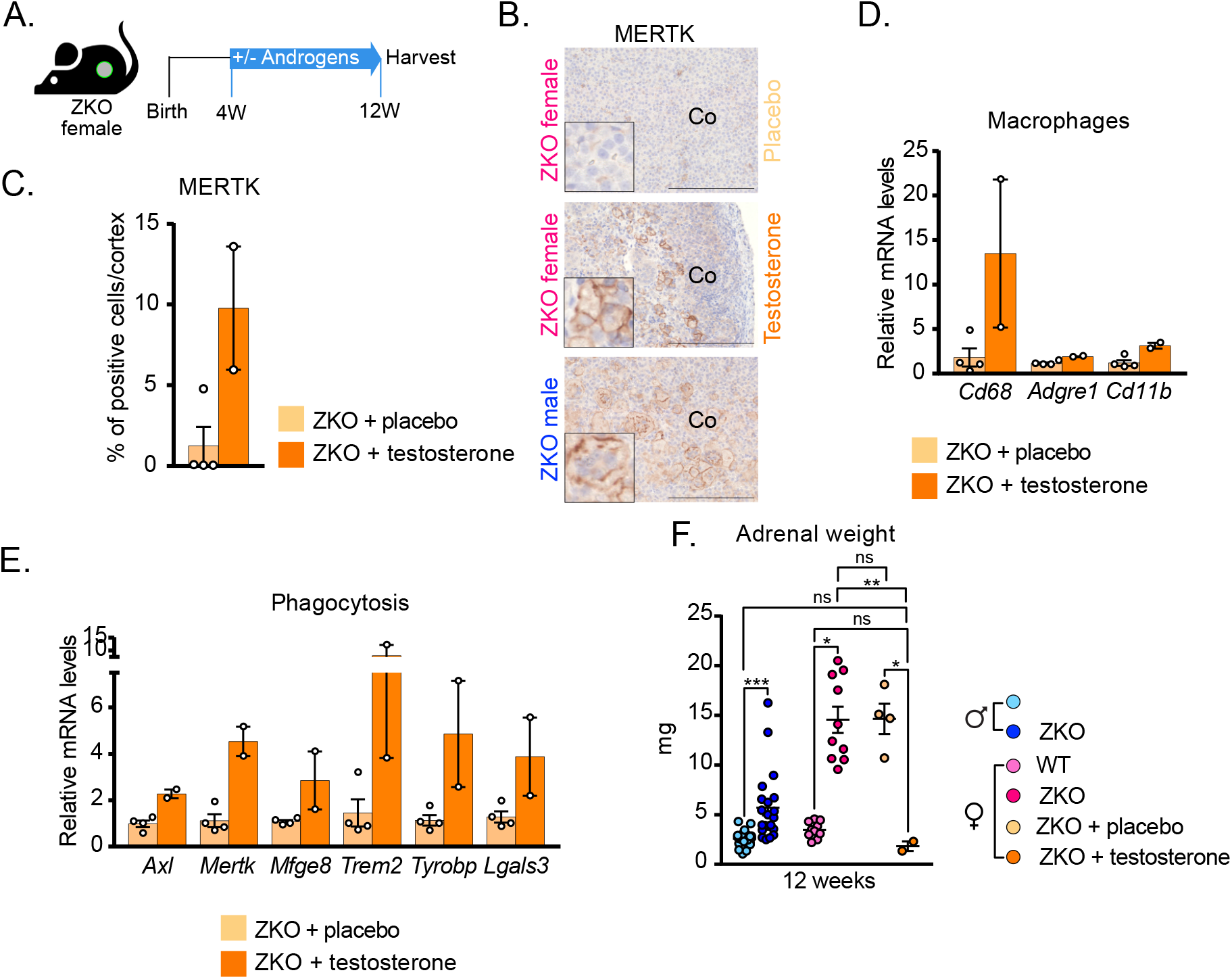
Androgens are sufficient to trigger early recruitment of phagocytic macrophages and regression of hyperplasia. **A-** Cartoon of the experimental setup. **B-** Immunohistochemical analysis of MERTK expression in 12-week-old placebo and testosterone-treated *Znrf3 cKO* females. An untreated 12-week-old *Znrf3 cKO* male was included as a reference. **C-** Quantification of the MERTK^+^ index as the ratio of MERTK-positive cells over total nuclei in the cortex of placebo and testosterone-treated females at 12 weeks. **D-** RTqPCR analysis of the expression of macrophages-related genes in placebo and testosterone-treated *Znrf3 cKO* female adrenals at 12 weeks. **E-** RTqPCR analysis of the expression of phagocytosis and macrophages fusion-associated genes in placebo and testosterone-treated *Znrf3 cKO* female adrenals at 12 weeks. **F-** Adrenal weights from placebo and testosterone-treated 12-week-old *Znrf3 cKO* females. 12-week-old untreated control males/females *and Znrf3 cKO* males/females from Fig 1A&E were included as a reference. Co : cortex; Scale bar = 200 μm. Graphs represent mean +/- SEM. Statistical analyses in D and E were conducted by Mann-Whitney tests. In F, statistical analyses were conducted by one-way ANOVA followed by a Kruskal-Wallis *post-hoc* test. ns: not significant; * p<0.05; ** p<0.01; *** p<0.001.

### Recruitment of phagocytic macrophages in male *Znrf3 cKO* mice is associated with sexually dimorphic induction of senescence

Recruitment of myeloid cells to preneoplastic lesions has been associated with induction of senescence ^7,32^. To evaluate a potential role of senescence in the sexually dimorphic recruitment of phagocytes in the adrenal cortex of *Znrf3 cKO* mice, we evaluated enrichment of senescence-associated signatures in males and females from 4 to 12 weeks. Whereas most of these signatures were significantly enriched in *Znrf3 cKO* males at 6 and 12 weeks, there was no or negative enrichment in females (Fig 6A). This suggested that ablation of *Znrf3* resulted in male-specific induction of senescence. To further evaluate this hypothesis, we first analysed expression of the cell-cycle inhibitor p21. In these experiments, steroidogenic cells were labelled by GFP, which was expressed by the mTmG locus following SF-1:Cre-mediated recombination. Consistent with induction of senescence, there was a significant increase in p21 labelling-index within GFP^+^ steroidogenic cells in *Znrf3 cKO* males at 4 weeks (Fig 6B and S6A-B). Levels of P21^+^ cells accumulation returned to normal at 6 weeks in *Znrf3 cKO* males and were significantly reduced at 12 weeks, consistent with phagocytosis of senescent cells (Fig 6B). Surprisingly, a significant increase in P21 labelling was also observed in *Znrf3 cKO* females at 4 weeks and maintained up to 12 weeks (Fig 6B and S6B). This suggested that cell cycle was arrested in both males and females, following *Znrf3* ablation. To further assess induction of senescence, we analysed activity of the prototypic senescence-associated acidic β-galactosidase (SA-bGal). This showed a few positive cells in the subcapsular area and at the cortical-medullary junction in control males and females, which was further increased in control females at 12 weeks. This suggested that spontaneous senescence was taking place in these regions (Fig 6C). Strikingly, SA-bGal staining was increased within the inner cortex of male *Znrf3 cKO* mice at 6 weeks and to a lesser extent at 12 weeks, consistent with phagocytic clearance of senescent cells in male adrenals (Fig 6C). In contrast, there was no increase in SA-bGal staining in *Znrf3 cKO* females, which displayed a similar pattern to controls (Fig 6C). This suggested that although proliferation was arrested in both males and females, senescence was only induced in male *Znrf3 cKO* adrenals. To further confirm this, we evaluated expression of a senescence associated secretory phenotype (SASP) in our RNA sequencing data. This analysis showed that the 23 SASP-coding genes that were significantly deregulated in 12-week-old *Znrf3 cKO* male adrenals were not deregulated in females (Fig 6D), suggesting that establishment of a SASP was male-specific. This was further confirmed by RTqPCR analyses showing significant up-regulation of *Mmp12* and *Il1a* at 6 weeks and *Cxcl2, Mmp12, Il1a* and *Tnfrsf1b* at 12 weeks in male, but not female adrenals (Fig 6E & Fig S6C), as well as male-specific enrichment of gene sets for NFkB signalling, which plays a key role in SASP induction^44,45^ (Fig S6D). Interestingly, *Znrf3 cKO* female mice that received testosterone (Fig 5) also showed induction of senescence-associated β-galactosidase after 1 week of treatment (from 4 to 5 weeks, Fig 6F). This was associated with up-regulation of the SASP factors *Mmp12, Il1a* and *Tnfrsf1b* (Fig 6G), which was concomitant with recruitment of MERTK^high^ macrophages and regression of hyperplasia after testosterone treatment from 4 to 12 weeks (Fig 5). This suggested that testosterone played a key role in senescence induction, which in turn, allowed recruitment of macrophages through SASP factors. Consistent with this hypothesis, F4/80-positive macrophages were found in very close proximity to SA-bGalactosidase- and GFP-positive steroidogenic cells in the adrenal cortex of male *Znrf3 cKO* mice at 6 weeks (Fig 6H).

**Figure 6.**
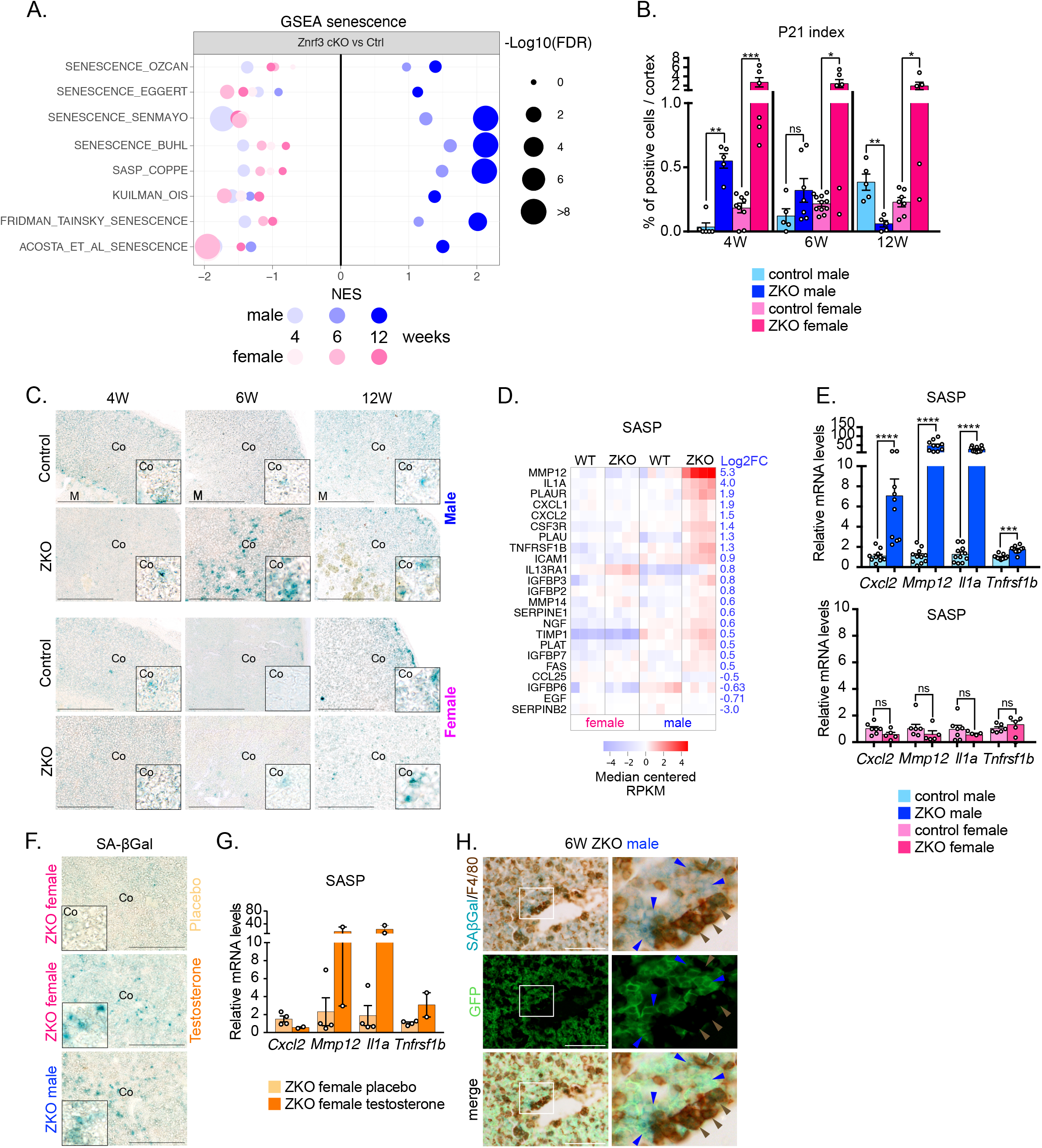
Recruitment of phagocytic macrophages in male *Znrf3 cKO* mice is associated with sexually dimorphic induction of senescence. **A-** Gene Set Enrichment Analysis (GSEA) of gene expression data (RNA sequencing) from 4, 6 and 12 control and *Znrf3 cKO* male and female adrenals. The plot represents enrichment of senescence-associated gene sets in *Znrf3 cKO* compared with controls (sex matched). **B-** Quantification of the P21^+^ index as the ratio of P21-positive cells over total nuclei in the cortex of male and female control and *Znrf3 cKO* mice from 4 to 12 weeks. **C-** Detection of the senescence associated acidic b-galactosidase activity on frozen tissue sections from male and female control and *Znrf3 cKO* mice at 4, 6and 12 weeks. Sections were counterstained with hematoxylin. **D-** Heatmap showing expression of senescence associated secretory phenotype (SASP) genes in 12-week-old male and female control and *Znrf3 cKO* adrenals. Genes were selected on the basis of significant deregulation in 12-week-old male *Znrf3 cKO* adrenals (FDR<0.1) and sorted by Log2 fold-change. **E-** RTqPCR analysis of the expression of SASP genes in control and *Znrf3 cKO* males (E - top panel) control and *Znrf3 cKO* females (E-bottom panel). **F-** Detection of SAb-Galactosidase activity in the adrenals of Znrf3 cKO females that received placebo or testosterone treatment from 4 to 5 weeks. An untreated 6-week-old *Znrf3 cKO* male was included as a reference. **G-** RTqPCR analysis of the expression of SASP genes in placebo and testosterone-treated *Znrf3 cKO* females from Fig 5. **H-** Immunohistochemical analysis of GFP (marking SF-1:Cre-mediated recombination of mTmG in steroidogenic cells), F4/80 and SAbGalactosidase activity. Right panels show a high-magnification crop of the area delineated in white in left panels. Blue arrowheads show senescent GFP^+^ cells; brown arrowheads show F4/80^+^ macrophages. Co: cortex; Scale bar = 200 μm (C-F); 100 μm (H). Graphs represent mean +/- SEM. Statistical analyses in B and E were conducted by Mann-Whitney tests. ns: not significant; * p<0.05; ** p<0.01; *** p<0.001; **** p<0.0001.

Altogether, these data strongly suggest that male-specific androgen-driven induction of senescence and SASP, results in recruitment, activation and fusion of highly efficient phagocytes that prevent tumour progression in male *Znrf3 cKO* mice.

### Aggressive tumourigenesis is associated with infiltration of non-phagocytic macrophages in female adrenals

To further gain insight into the role of macrophages at late stages of tumourigenesis, we evaluated infiltration of macrophages in 78-week-old adrenal lesions in both male and female mice. At this stage, male *Znrf3 cKO* adrenals were still infiltrated by IBA-1-positive macrophages that were scattered throughout the cortex (Fig 7A). However, quantification of the IBA-1 index showed that in contrast with earlier stages, infiltration was equivalent to control males (Fig 7B). In female *Znrf3 cKO*, IBA-1-positive infiltration was somewhat heterogenous within the tissue, with areas of high infiltration and zones that were almost devoid of macrophages (Fig 7A). There was also interindividual heterogeneity. Indeed, some tumours were still infiltrated at levels comparable to controls, whereas others showed much less IBA-1-positive cells or virtually no macrophages (Fig 7A & B). There was no overall difference between indolent and aggressive (metastatic) tumours with respect to IBA-1 index (Fig 7B).

**Figure 7.**
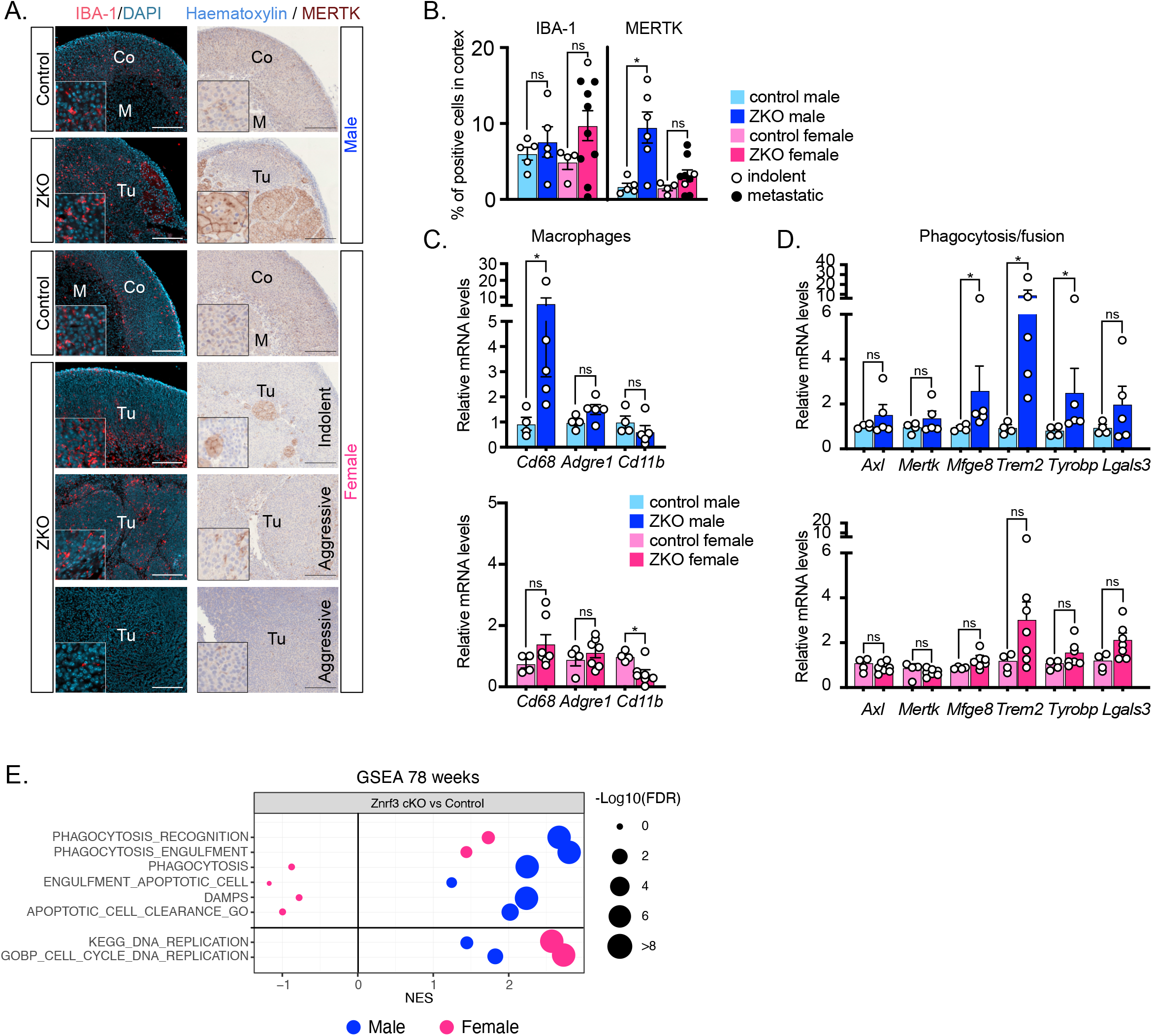
Aggressive tumourigenesis is associated with infiltration of non-phagocytic macrophages in female adrenals. **A-** Immunohistochemical analysis of IBA-1 and MERTK expression in control males/females and *Znrf3 cKO* males/females at 78 weeks. For *Znrf3 cKO* females, the panels represent indolent tumours (no metastases) and aggressive tumours with or without macrophages infiltration. **B-** Quantification of the IBA-1^+^ and MERTK^+^ index as the ratio of IBA-1 (left) or MERTK-positive (right) cells over total nuclei in the cortex of male and female control and *Znrf3 cKO* mice at 78 weeks. Values for primary tumours associated with metastases are shown as black dots. **C-** RTqPCR analysis of the expression of macrophages-related genes in control and *Znrf3 cKO* males (top panel) and control and *Znrf3 cKO* females (bottom panel) at 78 weeks. **D-** RTqPCR analysis of the expression of phagocytosis and macrophages fusion-associated genes in control and *Znrf3 cKO* males (top panel) and control and *Znrf3 cKO* females (bottom panel) at 78 weeks. **E-** Gene Set Enrichment Analysis (GSEA) of gene expression data (RNA sequencing) from 78 weeks control and *Znrf3 cKO* male and female adrenals. The plot represents enrichment of phagocytosis-associated gene sets and DNA replication-associated gene sets in *Znrf3 cKO* compared with controls (sex matched). Co: cortex; Tu: tumour; Scale bar = 200 μm. Graphs represent mean +/- SEM. Statistical analyses in B and E were conducted by Mann-Whitney tests. ns: not significant; * p<0.05.

However, macrophages exclusion was only observed in a subset of 2/10 aggressive tumours (Fig 7A & B). Consistent with IHC analyses, accumulation of mRNA encoding macrophages markers were unaltered (*Cd68*, *Adgre1*) or decreased (*Cd11b*) in female *Znrf3 cKO* compared to controls (Fig 7C). Interestingly, although accumulation of *Adgre1* and *Cd11b* mRNA was unaltered, *Cd68* was still strongly accumulated in male *Znrf3 cKO* adrenals (Fig 7C). Since we showed high expression of CD68 in fused macrophages at earlier stages (Fig 2G), this suggested that active phagocytes may still be accumulating in male KO adrenals at 78 weeks. Consistent with this idea, there were still large numbers of MERTK^high^ fused macrophages in 78-week-old *Znrf3 cKO* male adrenals (Fig 7A &B), which was correlated with overexpression of *Mfge8, Trem2* and *Tyrobp* in qPCR (Fig 7D). In contrast, female *Znrf3 cKO* adrenals showed scarce infiltration of MERTK^high^ fused macrophages, although a few of them could still be observed in indolent tumours (Fig 7A & B). Consistent, with these observations, there was no deregulation of phagocytosis/fusion markers in *Znrf3 cKO* female adrenals at this stage (Fig 7D). Interestingly, gene set enrichment analyses showed strong enrichment of phagocytosis-associated signatures, but no enrichment of DNA proliferation / cell-cycle pathways in male *Znrf3 cKO* RNA sequencing data at 78 weeks (Fig 7E). In contrast, female knockouts showed high enrichment of proliferation signatures, but no enrichment of phagocytosis (Fig 7E).

Altogether, these data strongly suggest that even though macrophages are still present within tumour tissues at 78 weeks in both males and females, the lack of phagocytic activity is associated with aggressive tumour progression in females.

### Phagocytic macrophages signatures are prominent in male ACC patients and associated with better prognosis

To further evaluate the role of macrophages in ACC progression, we evaluated their infiltration within human ACC. For this, we used RNA sequencing data from the TCGA consortium (79 sequenced ACCs) and evaluated expression of a 10-gene signature (based on single cell RNA sequencing data from mouse adrenals) as a proxy to general macrophages infiltration. Interestingly, tumours of the good prognosis group, defined as C1B, had significantly higher expression of the macrophages signature than tumours of the bad prognosis C1A group (Fig 8A). Consistent with our data showing similar infiltration of IBA-1^+^ macrophages in male and female *Znrf3 cKO* adrenals at 78 weeks (Fig 7B), there was no difference in the general macrophage signature expression between ACC in men (n=31) and women (n=48) (Fig 8B). However, a three gene phagocytic macrophage signature (*CD68, TREM2, TYROBP*) was significantly expressed at higher levels in men (Fig 8C), and in the good prognosis C1B group of ACC (Fig 8D). High expression of the phagocytic signature (above median) was also associated with better survival, compared with low expression (below median) in Kaplan-Meier analysis (Fig 8E). Altogether, this suggested that infiltration with phagocytic macrophages was more frequent in men than in women and associated with better prognosis.

**Figure 8.**
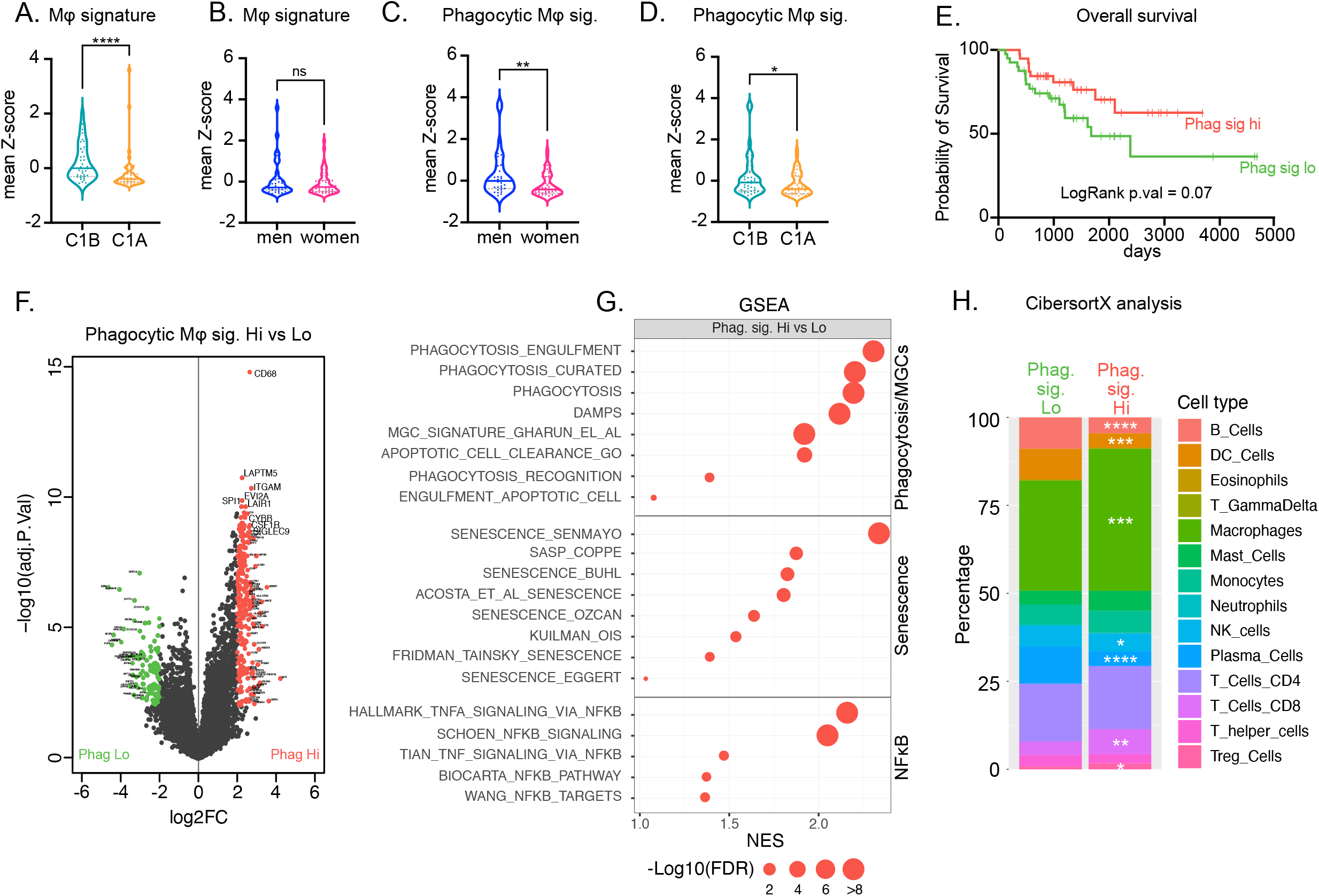
Phagocytic macrophages signatures are prominent in male ACC patients and associated with better prognosis. **A-** Expression of a global macrophage gene signature in ACC patients from the TCGA program, dichotomised as patients with good (C1B) and poor (C1A) prognosis. **B-** Expression of a global macrophage gene signature in ACC patients from the TCGA program, dichotomised as men and women. **C-** Expression of a phagocytic macrophage gene signature in ACC patients from the TCGA program, dichotomised as men and women. **D-** Expression of a phagocytic macrophage gene signature in ACC patients from the TCGA program, dichotomised as patients with good (C1B) and poor (C1A) prognosis. **E-** Survival analysis of patients of the TCGA program dichotomised as patients with high (red) or low (green) expression of the phagocytic signature. **F-** Volcano plot displaying differential gene expression between patients with high and low expression of the phagocytic signature. Red dots represent genes with a Log2 fold-change> 2 and FDR<0.01. Green dots represent genes with Log2 fold-change<-2 and FDR<0.01. **G-** GSEA of phagocytosis, senescence and NFkB-associated gene sets in patients with high expression of the phagocytic signature, compared with patients with low expression of the signature.

Detailed analysis of RNA sequencing data identified 365 genes that were significantly deregulated (FDR <0.01, abs(Log2FC>2)) between the groups of high and low expression of the phagocytic signature (Fig 8F). As expected, macrophages-associated genes such as *CD68*, *CSF1R, ITGAM, LAPTM5, CYBB* and *SIGLEC9* were up-regulated in phagocytic-high patients (Fig 8F). Gene set enrichment analyses confirmed enrichment of macrophages (Fig S7A) and phagocytosis signatures (Fig 8G). Consistent with data in our mouse models, phagocytic signatures were also associated with enrichment of senescence and NFkB signalling gene sets (Fig 8G), suggesting that these pathways may also play a role in phagocytic macrophages recruitment in ACC patients. Gene ontology analysis using the C5 GO database (MsigDB) showed that the top 35 positively enriched gene sets were all related with immune response and inflammation in patients with high expression of the phagocytic signature, suggesting that this subgroup was mounting a more profound immune response than patients with low expression of the signature (Fig S7B). Deconvolution of RNAseq data using Cibersort X showed that macrophages were the most prominent immune cell population in the two groups of patients, consistent with mouse adrenals (Fig 8H). Interestingly, it also showed that enrichment of macrophages signatures in the phagocytic-high subgroup of patients was associated with increased cytotoxic CD8^+^ T lymphocytes signatures (Fig 8H). However, this was also correlated with lower B cells, plasma cells and NK cells and higher T-regulatory cells infiltration (Fig 8H), suggesting that the phagocytic-high subgroup of patients had a broad alteration of the immune tumour microenvironment.

Altogether, these observations suggest that phagocytic macrophages, which are more prominent in male than female ACC patients, are associated with senescence, global innate and adaptive immune response and better prognosis.

## Discussion

Apart from reproductive tissues, cancers are generally more frequent and aggressive in men than in women, even after adjusting for known risk factors^1,2^. Although adrenocortical carcinoma is one of the rare exceptions to this rule, the mechanisms underlying the higher incidence and aggressiveness in women remain elusive. Here, we show that conditional deletion of *Znrf3* within steroidogenic cells of the adrenal cortex, results in sexually dimorphic development of full-fledged metastatic ACC in female mice over an 18-month time course, whereas the initial hyperplasia gradually regresses in males. By a combination of RNA sequencing, flow cytometry and immunohistochemical analyses, we show that *Znrf3 cKO* males efficiently recruit macrophages from early stages of preneoplastic transformation, following induction of senescence. We further show that these macrophages, which differentiate as potent phagocytes are required for clearance of preneoplastic cells. Although females also mount an innate immune response to preneoplastic transformation, it is delayed compared to males and never achieves efficient clearance of preneoplastic cells. This phenomenon is maintained up to 78 weeks, when indolent lesions in male *Znrf3 cKO* adrenals are still infiltrated with large amounts of phagocytic macrophages, as opposed to aggressive female tumours. Consistent with our findings in mice, we show that a phagocytic macrophage signature is more prominent in male than in female ACC patients, where it is associated with better prognosis. This strongly suggests that the sexual dimorphism of ACC may result from differential recruitment and activation of phagocytic tumour associated macrophages (TAMs), which prevent both tumour initiation and progression in the adrenal cortex.

This is in contrast with most data of the literature showing that TAMs are generally associated with tumour progression and poor prognosis in many cancers, even though they may initially prevent tumour initiation^7,32,46–48^. Plasticity and diversity of TAMs explain their divergent functions. The standard dual classification of macrophages postulates that M1 macrophages that differentiate in response to proinflammatory cytokines (*e.g*. interferons and tumour necrosis factors) are involved in anti-tumour activities, whereas M2 macrophages that differentiate in response to immunomodulatory signals (*e.g*. IL-4, IL-10 and TGF-β) are associated with tumour promotion ^49^. However, recent single cell RNA sequencing analyses of tumour infiltrating myeloid cells showed that M1 and M2 gene signatures were co-expressed in macrophage subsets from almost all cancer types^47^. Consistent with this idea, our RNA sequencing and flow cytometry analyses suggested that macrophages that accumulate in the adrenals of *Znrf3 cKO* males had mixed characteristics of the M1 and M2 phenotypes. Furthermore, we did not find evidence of overexpression of canonical tumoricidal macrophages markers such as the pro-inflammatory cytokines IL-1b, IL-2, IL-6, IL-12 and IL-23 (Fig 2B) or iNOS, which metabolizes arginine into the killer molecule nitric oxide (not shown). This suggests that the tumoricidal function of adrenal macrophages relies on alternative activities.

Consistent with this idea, we show a very strong increase in the phagocytic activity of macrophages in *Znrf3 cKO* male adrenals compared with their wild-type littermates and *Znrf3 cKO* females. This activity is associated with cytoplasmic accumulation of CD68 and high membrane expression of MERTK, TYROBP and TREM2, which play a central role in the phagocytic process ^35,38,40–42^. Although *MERTK* expression did not correlate with the presence of macrophages in ACC patients (not shown), we show that increased expression of the phagocytic *CD68/TREM2/TYROBP* signature is correlated with better prognosis, within the TCGA cohort. This strongly suggests that phagocytosis plays a central role in the tumoricidal activity of macrophages in ACC. Interestingly, scRNA sequencing in human and mouse colorectal cancer identified a population of C1QC^+^ TAMs, characterised by high levels of *C1QA/B/C, TREM2* and *MERTK* expression, which were associated with potential recruitment and activation of T cells, phagocytosis and better prognosis^47,50^. Although we did not analyse macrophages by single cell RNA sequencing in *Znrf3 cKO* adrenals, our bulk RNA sequencing data show strong up-regulation of all these markers (Fig 3B), which are mostly expressed by macrophages in scRNA seq datasets from wild-type mouse adrenals (Fig S8A). This strongly suggests that tumoricidal TAMs found in ACC may be related with the phagocytic C1QC^+^ TAMs identified in other cancers^47,50^.

Interestingly, we could find large numbers of macrophages (up to 15% of total cells in the tumour) in aggressive tumours in 78-week-old *Znrf3 cKO* female mice (Fig 7B), even though they were not differentiated as MERTK-hi active phagocytes. The presence of macrophages was further confirmed in ACC patients, where Cibersort X deconvolution suggested that they represented 31% of all immune cells, even in the tumours expressing low levels of the phagocytic signature (40% in phagocytic high tumours, Fig 8H). These data suggest that even in aggressive phagocytic-low lesions, macrophages may be reprogrammed to stimulate their tumoricidal potential. A large panel of molecules targeting macrophages is now available. Most of these pharmacological compounds or monoclonal antibodies aim at reducing macrophages infiltration within the tumour microenvironment, which our data suggest is not the best approach in ACC. However, strategies that stimulate tumoricidal activity and particularly phagocytosis of tumour cells by macrophages are currently being investigated^49^. Particularly interesting is the approach that aims at inhibiting the CD47 “don’t-eat-me” signal produced by cancer cells and/or the SIRPa receptor for CD47 on macrophages. In preclinical mouse models, this approach allowed stimulation of phagocytosis and tumour regression and also enhanced tumour antigen cross-presentation, resulting in adaptive immune responses^51,52^. It is currently evaluated in a number of cancers such as non-hodgkin lymphomas and acute myeloid leukaemia, where it results in good overall response rates in the absence of overt toxicities^49^. Another interesting approach aims at stimulating Toll-Like Receptor signalling (TLR) with TLR agonists, which stimulates macrophages tumoricidal activity and allows for secretion of IL-12 and TNF, which promotes a cytotoxic CD8^+^ T cell response. Agonists such as FDA-approved imiquimod (TLR7) or vidotilimod (TLR9) provide interesting responses in the context of basal cell carcinoma and metastatic melanoma, respectively^49^. Monophosphoryl lipid A (MPLA) a TLR4 agonist that is used as an FDA-approved vaccine adjuvant, was also demonstrated to trigger efficient innate and adaptive immune responses in association with IFN-γ, in the context of preclinical mouse models of breast and ovarian tumours^53^. One important factor that these therapeutic approaches will have to consider in the context of ACC, is the presence of high levels of glucocorticoids produced by adrenal steroidogenic cells, in particular within hormonally active tumours. Although glucocorticoids do not have the same detrimental impact on macrophages that they have on lymphocytes, they are generally associated with M2-like tolerogenic differentiation^54^. Therefore, therapeutics targeting macrophages in ACC, should probably consider combining macrophage activation with inhibition of glucocorticoid production or signalling, which would also favour recruitment of adaptive immune cells to the lesion. Availability of our clinically relevant mouse model will allow evaluation of these novel options.

An intriguing feature of the immune response following *Znrf3* deletion, is the formation of multinucleated giant cells (MGCs), resulting from the fusion of mononucleated macrophages. MGCs were first described in tuberculosis but are also present in sterile chronic inflammatory conditions and in response to macroscopic foreign bodies ^55,56^. Literature on their association with tumours is rather scarce. However, they have been observed at high frequency in squamous cell carcinomas of multiple tissues^57^ and papillary thyroid carcinomas^58^. They can either be correlated with good^57^ or poor prognosis^58^ depending on the tumour type and in vitro studies have associated MGCs with increased capacity for complement-mediated phagocytosis of large targets and amyloid deposits^55^. In *Znrf3 cKO* adrenals, MGCs are characterised by high levels of CD68, F4/80, MERTK, TYROBP and TREM2 expression and drastically reduced expression of IBA-1 (Fig 2G-I & Fig S4C-D). They appear early in male adrenals, almost concomitant with regression of hyperplasia and are maintained up to 78 weeks. In contrast, they only appear at late stages in females, are less frequent than in males and are mostly found associated with indolent non-metastatic lesions at 78 weeks. Although we did not analyse macrophages *in situ* in ACC samples from patients, we further observed enrichment of a gene signature associated with development of MGCs from monocyte progenitors^59^, within the high-phagocytosis group of patients. Altogether, this suggests that MGCs may play a role in clearing out neoplastic cells within both mouse and human tumours. Alternatively, their appearance may be a by-product of phagocytic clearance of neoplastic cells by mononuclear macrophages that would then fuse to form MGCs. This scenario could be facilitated by the high expression of TYROBP and TREM2 which have been shown to establish a fusion-competent state for macrophages^31^. Progressive cholesterol accumulation resulting from phagocytosis of cholesterol-laden steroidogenic cells, may also play a role in this phenomenon. Indeed, cholesterol is an essential and rate-limiting factor for formation of MGCs^59^ and oil-red-o staining of *Znrf3 cKO* male adrenals at 12 weeks showed a strong accumulation of lipids within MGCs (Fig S8B).

Another striking feature of the phenotype is the strong sexual dimorphism in immune response to neoplasia. It culminates with the robust and early recruitment of tumoricidal phagocytic macrophages in male mice, which results in regression of hyperplasia and prevents aggressive tumorigenesis, specifically in this gender. We further show that testosterone treatment of females from 4 to 12 weeks is sufficient to trigger a response which is comparable to males and results in regression of hyperplasia (Fig 5). This strongly suggests a role of male hormones in this phenomenon and raises the question of the underlying mechanisms. One possibility is an intrinsic sexual dimorphism of macrophages within the adrenal, which would result in differential responses to oncogenic transformation of steroidogenic cells. Indeed, recruitment, replenishment and activation mechanisms of macrophages have been shown to diverge between males and females, resulting in sexually dimorphic responses to infection and proinflammatory stimuli. However, in most instances, female macrophages are more responsive to stimuli, mount a more robust response and have higher phagocytic capacities than male macrophages^60–65^. This suggests that the stronger inflammatory response observed in male *Znrf3 cKO* adrenals may result from indirect effects of sex hormones. Consistent with this, single cell sequencing data suggest that the androgen receptor *Ar* is only expressed in a small subset of adrenal macrophages, characterized by lower expression of *Trem2* and *Mertk*, which is unlikely to represent the major population of macrophages in *Znrf3c KO* adrenals (Fig S8A&C). Our data showing a strong association between induction of SASP and recruitment of macrophages suggest that androgens may stimulate the tumoricidal response by inducing release of senescence associated cytokines by *Znrf3 cKO* cells (Fig 6). In line with this hypothesis, AR activation was shown to induce p53-independent senescence in prostate cancer cells^66,67^ and a short-term testosterone treatment was sufficient to induce SA-bgalactosidase activity in female *Znrf3 cKO* adrenals (Fig 6). This raises the question of the links between *Znrf3* inactivation, AR signalling and senescence induction. One may speculate that the recently documented sexual dimorphism in cortical cell proliferation, renewal and progenitor populations ^8,9^ may result in sexually dimorphic response to *Znrf3* inactivation. In this context, the hyperproliferation observed in both male and female *Znrf3 cKO* adrenals may result in faster exhaustion of progenitor pools in males and subsequent induction of senescence. However, the rapid induction of SA-bgalactosidase in testosterone-treated *Znrf3 cKO* females suggests that this is an unlikely scenario. Alternatively, these findings may reflect a novel function of ZNRF3 in the control of cellular homeostasis. Interestingly, although we were able to show a mild induction of *Axin2* accumulation in *Znrf3 cKO* adrenals by RNA in-situ hybridization^29^, analysis of our RNA sequencing data did not show evidence of canonical WNT signalling induction in either male or female knockouts (Fig S8D), when compared with a previously published model of constitutive β-catenin activation^68^. This suggests that the impact of *Znrf3* inactivation on senescence induction may involve WNT-independent mechanisms. Interestingly, inactivation of *Znrf3* and its homologue *Rnf43* in hepatocytes results in hyperplasia, followed by cell death and senescence induction. This is associated with deregulation of lipid and phospholipid metabolism through both canonical and non-canonical WNT pathway activation^69^. Whether similar mechanisms resulting in accumulation of toxic lipids are involved in senescence induction in the adrenal cortex, remains to be investigated.

In conclusion, we describe a novel interaction between tumour suppressor inactivation, senescence induction and recruitment of tumoricidal macrophages, which results in sexually dimorphic adrenal cancer development. This provides novel insight into the strong gender bias of this particularly aggressive cancer and may help develop novel macrophage-centric therapeutic approaches.

## Acknowledgments

We thank Sandrine Plantade, Khirredine Ouchen and Philippe Mazuel for animal care and Dr Laura Bousset (LMS, London) for fruitful discussions. The TREM2 antibody was supplied by the Haass Lab at Ludwig Maximilians University Munich, specifically by Alice Suelzen and Dr Kai Schlepckow. Single cell RNA sequencing data from mouse adrenals were kindly provided by Dr Cynthia Andoniadou and Val Yianni (King’s College London). Protocol for mouse adrenal tissue dissociation for flow cytometry and flow cytometry parameters were kindly provided by Dr Marc Bajénoff and Dr Mitchell Bijnen (CIML, Marseille). This work was supported by grants from Worldwide Cancer Research (#16-1052), la Ligue Nationale Contre le Cancer (“Equipe Labellisée Ligue” and individual grant to JJW), Fondation ARC (individual grant to JO) and Agence Nationale pour la Recherche (ANR-21-CE14-0044-ADREMAC).

## Authors contributions

JJW performed experiments, analysed data, prepared the manuscript

JO performed experiments, analysed data, prepared the manuscript

DGG performed experiments, analysed data, prepared the manuscript

CL performed experiments

RG conceived experiments

FRB performed experiments

DD performed experiments

CDS performed experiments

ISB performed experiments

JCP performed experiments

YR analysed data

AL performed experiments

IT conceived experiments, discussed findings

AMLM conceived experiments, discussed findings

AM conceived experiments, discussed findings

PV conceived experiments, performed experiments, analysed data, wrote the manuscript

## Materials and Methods

### Mice

All experiments with mice were in accordance with protocols approved by the Auvergne Ethics Committee (CEMEAA). They were conducted in agreement with international standards for animal welfare in order to minimize animal suffering. *Znrf3 cKO* mice (ZKO) were generated by mating *Znrf3^fl/fl^* mice^27^ with *SF1-Cre^high^* mice^70^. Mice were bred and maintained on a C57Bl/6 genetic background. Mice were euthanized by decapitation at the end of experiments and blood collected in vacuum blood collection tubes (VFD053STK, Terumo). Adrenals were extracted, cleaned of excess fat, weighed and immediately fixed in 4%paraformaldehyde or stored at −80 °C. Littermate control animals were used in all experiments.

### Immunohistology

Adrenals were fixed in 4% paraformaldehyde overnight at 4°C, then washed two times in PBS. For the paraffin embedding, adrenals were dehydrated through an ethanol gradient. Then they were incubated for 2 h in Histoclear (HS200; National Diagnostics, Fisher Scientific, Illkirch, France) and embedded in paraffin. For frozen sections, adrenals were successively placed into 10% and 15% PBS-sucrose solutions for 20 minutes, then 20% PBS-sucrose solution for 1 hour, and in 50/50 OCT-Sucrose 20% solution overnight. Lastly, they were embedded in pure OCT solution and stored at −80°C. Paraffin and OCT samples were cut into 5 & 10μm sections, respectively. Haematoxylin/eosin staining was performed with a Microm HMS70 automated processor (Microm Microtech, Francheville, France), according to standard procedures. Antibody information, dilutions and unmasking conditions are listed in Supplementary Table 1. Notably, the TREM2 antibody ^71^, was supplied from the Haass Lab at Ludwig Maximilians University Munich. After deparaffinization with Histoclear and rehydration in decreasing ethanol gradients, unmasking was performed by boiling slides for 20 min in the appropriate unmasking solution. Next, endogenous peroxidases were inactivated by incubating slides with 0.3% hydrogen peroxide for 30 min at room temperature. After blocking for 1h, slides were incubated overnight at room temperature with primary antibodies at the indicated concentrations (Supplementary Table 1). Primary antibodies were detected with appropriate species polymers (ImmPress Polymer Detection Kit, Vector Laboratories). Polymer-coupled HRP activity was then detected with either Novared (SK-4800, Vector Laboratories) for brightfield images or TSA-Alexa-coupled fluorochromes for fluorescence (Thermo Fisher, Alexa_488 B40953, Alexa_555 B40955, Alexa_647 B40958). For double-immunohistochemistry experiments, HRP was inactivated by incubation with 0.02% HCl for 20 min after detection of the first antibody to avoid cross-reaction. Nuclei were counterstained with Haematoxylin for brightfield images or Hoechst for fluorescence (Thermo Fisher 33342). Slides were mounted using a 50/50 PBS-Glycerol solution. Images were acquired with a Zeiss AxioImager with Apotome2 or Zeiss Axioscan Z1 slide scanner. Images were minimally processed for global levels and white balance using Affinity Photo® and Affinity Designer®. Image settings and processing were identical across genotypes.

Quantifications were performed on scanned whole adrenals (Axio Scan Zeiss scanner, 20× images) using the QuPath software version 0.3.1 (Bankhead et al 2017). Briefly, annotations were made of whole adrenals or just the adrenal cortex, and the positive cell detection feature was used to identify positive cells. The threshold for identifying positive cells was set to avoid quantification of background on each image.

For quantification of phagocytosis, confocal images were acquired on a Zeiss LSM 800 Airyscan confocal microscope with 40X magnification. Phagocytic events were identified and counted as the presence of steroidogenic cell markers (3bHSD or SF-1) within the boundaries of macrophages, defined by IBA-1 or MERTK staining. This was evaluated by a single operator, by manually scanning through Z-stacks of ten 40X images per adrenal. The operator was blinded to the genotype.

### Senescence-associated beta-galactosidase (SA-β-galactosidase) staining

SA-β-galactosidase staining was conducted following the protocol of Debacq et al (4) on frozen adrenal 10μm-sections. After drying for 15 min under a vacuum, the sections were rehydrated with PBS and then incubated overnight at 37°C in a humid atmosphere in a pH 6.0 staining solution composed of 7.4mM citric acid, 25.3mM dibasic sodium phosphate, 5mM K_4_[Fe(CN)_6_], 5mM K_3_[Fe(CN)_6_], 150mM sodium chloride, 2mM MgCl_2_ and 1mg/mL X-gal. Slides were mounted using a 50/50 PBS-Glycerol solution and imaged on a Zeiss ApoTome microscope with an AxioCam MRm camera and/or a Axio Scan Zeiss scanner.

### Testosterone supplementation experiment

Testosterone or placebo implants were placed under gas anesthesia, in the interscapular region of 4-week-old *Znrf3 cKO* female mice for 60 days. These testosterone implants (T-M/60 Belma) are designed to release daily doses of testosterone (from 51.9 to 154.5 μg/24hr for plasma concentrations of 0.9-3.7 ng/ml) to produce physiological plasma concentrations in mice.

### Pexidartinib experiment

Chow was purchased from SAFE Nutrition Services (Augy, France). Male Control & ZKO mice were fed either control chow (E8220A01R 00000 v0025 A04 Pur) or pexidartinib chow (E8220A01R 00000 v0398 A04 +0.29g/kg Pexidartinib) from 3-12 weeks of age. Pexidartinib (HY16749) was purchased from MedChemExpress and incorporated in the chow by SAFE Nutrition Services. Chow was replaced every 3-4 days, renewed weekly and stored at 4°C when not in use.

### FACS

Adrenals were harvested and excess fat was removed under a dissecting microscope. Adrenals were immediately placed into 900μL of digestion medium (Supplementary Table 2) and placed on ice until the end of the harvest. Adrenals were digested by incubating with a thermomixer set at 37°C – 900 rpm- for 37 min, stopping to pipette up and down at 10, 20, 30, 35, & 37min. Digested samples were filtered through 100μm nylon mesh and centrifuged at 400g for 5 min at 4°C. Cells were resuspended in wash buffer (PBS – EDTA 2.5mM – DNAse 100μg/ml – BSA 0.5%)) and stained appropriately. Cells were stained with Fixable Near-IR live/dead stain (L34975, Invitrogen) for 30min at room temperature (RT), blocked with CD16/CD32 & TrueStain (426102, BioLegend) for 15 min at RT, and stained with the appropriate antibody panel for 20 min at RT (Supplementary Table 3). All staining/blocking steps were preceded and followed by wash steps which included centrifugation at 200g for 4 min, followed by resuspension of the pellet with either wash buffer or the appropriate solution. Cells were immediately analyzed on the Attune NxT Flow Cytometer (Reference: A24858). Detailed analyses of the results were done using FlowJo® software.

### Reverse-transcription quantitative PCR

Adrenals were flash-frozen and stored at −80°C post-harvest. RNAs were extracted using the Macherey-Nagel Nucleospin RNA kit (REF #740955.250). After reverse transcription of 500ng of total RNAs, cDNAs were diluted 1/10 and PCR reactions were conducted using SYBR qPCR Premix Ex Taq II Tli RNase H+ (TAKRR820W, Takara). Primers can be found in Supplementary Table 4. Relative expression was calculated using the 2^-ΔΔCT method.

### RNA sequencing for gene expression analysis

#### Library preparation and sequencing

RNA sequencing was performed by the GenomEast platform, a member of the ‘France Genomique’ consortium (ANR-10-INBS-0009). Library preparation was performed using TruSeq Stranded mRNA Reference Guide - PN 1000000040498. RNA-Seq libraries were generated from 300 ng of total RNA using TruSeq Stranded mRNA Library Prep Kit and IDT for Illumina - TruSeq RNA UD Indexes (96 Indexes, 96 Samples) (Illumina, San Diego, USA), according to manufacturer’s instructions. Briefly, following purification with poly-T oligo attached magnetic beads, the mRNA was fragmented using divalent cations at 94oC for 2 minutes. The cleaved RNA fragments were copied into first strand cDNA using reverse transcriptase and random primers. Strand specificity was achieved by replacing dTTP with dUTP during second strand cDNA synthesis using DNA Polymerase I and RNase H. Following addition of a single ‘A’ base and subsequent ligation of the adapter on double stranded cDNA fragments, the products were purified and enriched with PCR (30 sec at 98°C; [10 sec at 98°C, 30 sec at 60°C, 30 sec at 72°C] x 12 cycles; 5 min at 72°C) to create the cDNA library. Surplus PCR primers were further removed by purification using SPRI select beads (Beckman-Coulter, Villepinte, France) and the final cDNA libraries were checked for quality and quantified using capillary electrophoresis. Libraries were sequenced on an Illumina HiSeq 4000 sequencer as single read 50 base reads. Image analysis and base calling were performed using RTA version 2.7.7 and bcl2fastq version 2.20.0.422.

#### Genome mapping and differential gene expression analyses

Reads were filtered and trimmed to remove adapter-derived or low-quality bases using cutadapt v 3.2 and checked again with FASTQC v 0.11.7. Illumina reads were aligned to Mouse reference genome (mm10) with Hisat2 v 2.2.1. Read counts were generated for each annotated gene using R function “SummarizeOverlaps()” and RPKM were calculated for each gene. Differential expression analysis with multiple testing correction was conducted using the R Bioconductor DESeq2 package v 1.34.0.

#### Generation of heatmaps

Heatmaps to represent differential gene expression were generated with the *Biobase* and *gplots* packages in R. They represent median centered RPKM levels. Genes are either sorted by Log2 fold-change or by unsupervised clustering.

### Reanalysis of single cell sequencing of adult mouse adrenals

The Seurat R package^72^ was used to perform clustering analysis of single-cell data from Lopez et al. ^73^, available in the Gene Expression Omnibus GSE161751 (control adrenals from 10 week-old male mice). Raw sequencing data and annotated gene–barcode matrices were used for the input. Cells with more than 20 genes and genes expressed in more than 3 cells were selected for further analysis. After studying the distribution of count depth, number of genes, and mitochondrial read fraction, low-quality cells with less than 1000 counts, less than 400 genes detected, and percentage of mitochondrial gene counts higher than 25% were removed. Gene expression in each cell was then normalized by the total number of counts in the cell, multiplied by 10000 to get counts per 10000 (TP10K) and log-transformed to report gene expression as *E* = log(TP10K +1).

The top 2,000 highly variable genes with a z-score cutoff of 0.5 were then centred and scaled to have a mean of zero and standard deviation of one, and used as inputs for initial principal component analysis (PCA). The number of principal components (PCs) was chosen according to the PCElbowPlot function and JackStrawPlot function. Next, the Louvain algorithm implemented in Seurat was used to iteratively group cells together, with the goal of optimizing the standard modularity function. The resolution parameter for clustering was set at *r* = 1. The default Wilcoxon rank-sum test was used by running FindAllMarkers function in Seurat to find differentially expressed markers in each cluster. Finally, each cell type was annotated after extensive literature reading and searching for specific gene expression patterns. Violin plot representations were used for visualizing expression of the different markers.

### TCGA adrenocortical carcinoma data

TCGA gene expression and clinical ACC data were extracted from the TCGA database (The Cancer Genome Atlas). Distribution in the good (C1B) and poor prognosis (C1A) groups were previously defined, based on unsupervised clustering^26^. Expression data were standardized by the Relative Standard Error of the Mean (RSEM) algorithm and transformed into Log2 in order to refocus and symmetrize values’ distribution. The macrophage signature was defined as the mean expression (Z-score) of *CD74, CXCL2, CCL4, APOE, CCL3, CTSS, C1QA, C1QB, C1QC* and *AIF1*. These were found as highly up-regulated genes in macrophages in single cell RNA sequencing analyses of adult mouse adrenals^73^ (see above). For gene set enrichment analyses, TCGA ACC patients were dichotomized based on the expression of a phagocytic signature (mean expression (Z-score) of *TYROBP, TREM2* and *CD68*) with patients classified as high (expression above median) or low (expression below median). Differential gene expression between patients from the phagocytic high and phagocytic low groups was computed using the *limma* R package. The volcano plot representing differential expression between these two groups was generated in R with the *calibrate* library. Kaplan Meier analysis was conducted in GraphPad prism after dichotomization of patients according to expression of the phagocytic signature.

### Gene Set enrichment analyses

Gene set enrichment analyses were conducted on gene expression data from mouse models and TCGA ACC patients, using GSEA 4.1.0 with gene sets from the MsigDB and MGI Gene Ontology databases and with custom curated gene sets (Supplementary Table 5). Permutations were set to 1000 and were performed on gene sets. Phagocytosis gene sets were curated from an extensive search of the literature, including papers by Park et al.^74^, Lecoultre et al.^75^ and Janda et al^76^ and extracted from the MGI gene ontology database. Senescence gene sets were extracted from papers by Eggert et al.^32^, Kuilman et al.^77^, özcan et al.^78^, Acosta et al.^79^, Fridman et al.^80^, Coppé et al.^81,82^, Buhl et al.^83^ and Saul et al. ^84^. LM22 and ImmuCC gene sets were derived from gene expression signatures published in Newman et al.^85^ and Chen et al.^86^. To reduce the gene expression matrix into simple gene identifier lists for GSEA, genes in each of the lists were attributed to their cognate immune cell type based on their maximum of expression across all cell types. This resulted in gene signatures for each immune cell type that were then used in GSEA (Table). M0, M1 and M2 macrophages gene sets were further concatenated to result in global LM22 and ImmuCC macrophages gene sets. The mouse adrenal macrophages gene set was defined as the 100 most significantly upregulated genes within the two macrophages clusters (compared to all other clusters) in our reanalysis of the single-cell sequencing study of adult mouse adrenals by Lopez et al.^73^. The cytokine gene set was curated from an extensive search of the literature. NFκB and DNA replication gene sets were extracted from MsigDB C2, Hallmarks and C5 datasets.

GSEA output was either displayed as dot plots or enrichment curves. Dot plots represent the normalized enrichment score (NES) and FDR (size of dots defined as -log10(FDR)) and were drawn using the *ggplot2* library in R. Enrichment curves were drawn by feeding GSEA output to the *GSEA_replot* R function, developed by Thomas Kuilman (https://github.com/PeeperLab/Rtoolbox/blob/master/R/ReplotGSEA.R). Dot plots and enrichment curves were further processed in Affinity Designer® for colour matching and superimposition.

### CibersortX and mMCP analyses

CibersortX analyses were run on the CibersortX server (https://cibersortx.stanford.edu) using the LM22 matrix and a mixture file representing gene expression data in control and *Znrf3 cKO* adrenals at 4, 6 and 12 weeks or TCGA ACC patients’ data, dichotomized on the basis of high or low expression of the phagocytic signature (see TCGA adrenocortical carcinoma data). Output of CibersortX was then processed in R to concatenate subpopulations of macrophages, B cells, T-CD4, NK cells, DCs and mast cells. *Ggplot2* was then used to generate stacked bar plots representing the percentage of each immune cell population. Statistical analyses between genotypes or patients’ groups were computed using the Mann-Whitney test.

mMCP analyses were run in R using the mMCP counter package (https://github.com/cit-bioinfo/mMCP-counter), following instructions in Petitprez et al.^87^ Stacked bar plots were generated by *ggplot2* and statistical analyses conducted as above.

### Statistical analyses

Minimal sample size was set at n=3 allowing for detection of 40% increases/decreases with α=0.05, δ=0.4 and sd=1.0. Statistical analyses were conducted with R and GraphPad Prism 9. Normality of data was assessed using D’Agostino & Pearson normality test. Statistical analysis of normally distributed data was performed by two-tailed Student’s *t*-test (two groups) with or without Welch’s correction (as a function of variance) or one-way ANOVA (multiple groups), followed by Tukey’s multiple comparisons test. Analysis of non-normally distributed data was performed by two-tailed Mann & Whitney test (two groups) or Kruskal-Wallis test followed by Dunn’s multiple comparisons test (multiple groups). All bars represent the mean ± SEM.

## References

1. Dart, A. Sexual dimorphism in cancer. Nat. Rev. Cancer 20, 627 (2020).

2. Clocchiatti, A., Cora, E., Zhang, Y. & Dotto, G. P. Sexual dimorphism in cancer. Nat. Rev. Cancer 16, 330–339 (2016).

3. Audenet, F., Méjean, A., Chartier-Kastler, E. & Rouprêt, M. Adrenal tumours are more predominant in females regardless of their histological subtype: a review. World J. Urol. 31, 1037–1043 (2013).

4. Else, T. et al. Adrenocortical Carcinoma. Endocr. Rev. 35, 282–326 (2014).

5. Lyraki, R. & Schedl, A. The Sexually Dimorphic Adrenal Cortex: Implications for Adrenal Disease. Int. J. Mol. Sci. 22, 4889 (2021).

6. Ayala-Ramirez, M. et al. Adrenocortical carcinoma: clinical outcomes and prognosis of 330 patients at a tertiary care center. Eur. J. Endocrinol. 169, 891–899 (2013).

7. Kang, T.-W. et al. Senescence surveillance of pre-malignant hepatocytes limits liver cancer development. Nature 479, 547–551 (2011).

8. Dumontet, T. et al. PKA signaling drives reticularis differentiation and sexually dimorphic adrenal cortex renewal. JCI Insight 3, (2018).

9. Grabek, A. et al. The Adult Adrenal Cortex Undergoes Rapid Tissue Renewal in a Sex-Specific Manner. Cell Stem Cell 25, 290–296.e2 (2019).

10. Baudin, E. & Endocrine Tumor Board of Gustave Roussy. Adrenocortical carcinoma. Endocrinol. Metab. Clin. North Am. 44, 411–434 (2015).

11. Sada, A. et al. Comparison between functional and non-functional adrenocortical carcinoma. Surgery 167, 216–223 (2020).

12. Shariq, O. A. & McKenzie, T. J. Adrenocortical carcinoma: current state of the art, ongoing controversies, and future directions in diagnosis and treatment. Ther. Adv. Chronic Dis. 12, 20406223211033104 (2021).

13. Fassnacht, M. et al. Combination chemotherapy in advanced adrenocortical carcinoma. N. Engl. J. Med. 366, 2189–2197 (2012).

14. Lo Iacono, M. et al. Molecular Mechanisms of Mitotane Action in Adrenocortical Cancer Based on In Vitro Studies. Cancers 13, 5255 (2021).

15. Puglisi, S. et al. New perspectives for mitotane treatment of adrenocortical carcinoma. Best Pract. Res. Clin. Endocrinol. Metab. 34, 101415 (2020).

16. Terzolo, M. et al. Adjuvant mitotane treatment for adrenocortical carcinoma. N. Engl. J. Med. 356, 2372–2380 (2007).

17. Berruti, A. et al. Long-Term Outcomes of Adjuvant Mitotane Therapy in Patients With Radically Resected Adrenocortical Carcinoma. J. Clin. Endocrinol. Metab. 102, 1358–1365 (2017).

18. Calabrese, A. et al. Adjuvant mitotane therapy is beneficial in non-metastatic adrenocortical carcinoma at high risk of recurrence. Eur. J. Endocrinol. 180, 387–396 (2019).

19. Le Tourneau, C. et al. Avelumab in patients with previously treated metastatic adrenocortical carcinoma: phase 1b results from the JAVELIN solid tumor trial. J. Immunother. Cancer 6, 111 (2018).

20. Carneiro, B. A. et al. Nivolumab in Metastatic Adrenocortical Carcinoma: Results of a Phase 2 Trial. J. Clin. Endocrinol. Metab. 104, 6193–6200 (2019).

21. Habra, M. A. et al. Phase II clinical trial of pembrolizumab efficacy and safety in advanced adrenocortical carcinoma. J. Immunother. Cancer 7, 253 (2019).

22. Raj, N. et al. PD-1 Blockade in Advanced Adrenocortical Carcinoma. J. Clin. Oncol. Off. J. Am. Soc. Clin. Oncol. 38, 71–80 (2020).

23. Thorsson, V. et al. The Immune Landscape of Cancer. Immunity 51, 411–412 (2019).

24. Landwehr, L.-S. et al. Interplay between glucocorticoids and tumor-infiltrating lymphocytes on the prognosis of adrenocortical carcinoma. J. Immunother. Cancer 8, e000469 (2020).

25. Assié, G. et al. Integrated genomic characterization of adrenocortical carcinoma. Nat. Genet. (2014) doi:10.1038/ng.2953.

26. Zheng, S. et al. Comprehensive Pan-Genomic Characterization of Adrenocortical Carcinoma. Cancer Cell 29, 723–736 (2016).

27. Koo, B.-K. et al. Tumour suppressor RNF43 is a stem-cell E3 ligase that induces endocytosis of Wnt receptors. Nature 488, 665–669 (2012).

28. Hao, H.-X. et al. ZNRF3 promotes Wnt receptor turnover in an R-spondin-sensitive manner. Nature 485, 195–200 (2012).

29. Basham, K. J. et al. A ZNRF3-dependent Wnt/β-catenin signaling gradient is required for adrenal homeostasis. Genes Dev. 33, 209–220 (2019).

30. Lucas, M. et al. Massive inflammatory syndrome and lymphocytic immunodeficiency in KARAP/DAP12-transgenic mice. Eur. J. Immunol. 32, 2653–2663 (2002).

31. Helming, L. et al. Essential role of DAP12 signaling in macrophage programming into a fusion-competent state. Sci. Signal. 1, ra11 (2008).

32. Eggert, T. et al. Distinct Functions of Senescence-Associated Immune Responses in Liver Tumor Surveillance and Tumor Progression. Cancer Cell 30, 533–547 (2016).

33. Truman, L. A. et al. CX3CL1/fractalkine is released from apoptotic lymphocytes to stimulate macrophage chemotaxis. Blood 112, 5026–5036 (2008).

34. Yang, L. V., Radu, C. G., Wang, L., Riedinger, M. & Witte, O. N. Gi-independent macrophage chemotaxis to lysophosphatidylcholine via the immunoregulatory GPCR G2A. Blood 105, 1127–1134 (2005).

35. Lemke, G. How macrophages deal with death. Nat. Rev. Immunol. 19, 539–549 (2019).

36. Galvan, M. D., Greenlee-Wacker, M. C. & Bohlson, S. S. C1q and phagocytosis: the perfect complement to a good meal. J. Leukoc. Biol. 92, 489–497 (2012).

37. Chen, J. et al. SLAMF7 is critical for phagocytosis of haematopoietic tumour cells via Mac-1 integrin. Nature 544, 493–497 (2017).

38. Cockram, T. O. J., Dundee, J. M., Popescu, A. S. & Brown, G. C. The Phagocytic Code Regulating Phagocytosis of Mammalian Cells. Front. Immunol. 12, 2144 (2021).

39. Hanayama, R. et al. Autoimmune disease and impaired uptake of apoptotic cells in MFG-E8-deficient mice. Science 304, 1147–1150 (2004).

40. Atagi, Y. et al. Apolipoprotein E Is a Ligand for Triggering Receptor Expressed on Myeloid Cells 2 (TREM2). J. Biol. Chem. 290, 26043–26050 (2015).

41. Nugent, A. A. et al. TREM2 Regulates Microglial Cholesterol Metabolism upon Chronic Phagocytic Challenge. Neuron 105, 837–854.e9 (2020).

42. Lu, Q. et al. Tyro-3 family receptors are essential regulators of mammalian spermatogenesis. Nature 398, 723–728 (1999).

43. Caberoy, N. B., Alvarado, G., Bigcas, J.-L. & Li, W. Galectin-3 is a new MerTK-specific eat-me signal. J. Cell. Physiol. 227, 401–407 (2012).

44. Meyer, P. et al. A model of the onset of the senescence associated secretory phenotype after DNA damage induced senescence. PLoS Comput. Biol. 13, e1005741 (2017).

45. Salminen, A., Kauppinen, A. & Kaarniranta, K. Emerging role of NF-κB signaling in the induction of senescence-associated secretory phenotype (SASP). Cell. Signal. 24, 835–845 (2012).

46. Vesely, M. D., Kershaw, M. H., Schreiber, R. D. & Smyth, M. J. Natural innate and adaptive immunity to cancer. Annu. Rev. Immunol. 29, 235–271 (2011).

47. Cheng, S. et al. A pan-cancer single-cell transcriptional atlas of tumor infiltrating myeloid cells. Cell 184, 792–809.e23 (2021).

48. Xue, W. et al. Senescence and tumour clearance is triggered by p53 restoration in murine liver carcinomas. Nature 445, 656–660 (2007).

49. Pittet, M. J., Michielin, O. & Migliorini, D. Clinical relevance of tumour-associated macrophages. Nat. Rev. Clin. Oncol. (2022) doi:10.1038/s41571-022-00620-6.

50. Zhang, L. et al. Single-Cell Analyses Inform Mechanisms of Myeloid-Targeted Therapies in Colon Cancer. Cell 181, 442–459.e29 (2020).

51. Weiskopf, K. et al. CD47-blocking immunotherapies stimulate macrophage-mediated destruction of small-cell lung cancer. J. Clin. Invest. 126, 2610–2620 (2016).

52. von Roemeling, C. A. et al. Therapeutic modulation of phagocytosis in glioblastoma can activate both innate and adaptive antitumour immunity. Nat. Commun. 11, 1508 (2020).

53. Sun, L. et al. Activating a collaborative innate-adaptive immune response to control metastasis. Cancer Cell 39, 1361–1374.e9 (2021).

54. Diaz-Jimenez, D., Kolb, J. P. & Cidlowski, J. A. Glucocorticoids as Regulators of Macrophage-Mediated Tissue Homeostasis. Front. Immunol. 12, 669891 (2021).

55. Milde, R. et al. Multinucleated Giant Cells Are Specialized for Complement-Mediated Phagocytosis and Large Target Destruction. Cell Rep. 13, 1937–1948 (2015).

56. Helming, L. & Gordon, S. Molecular mediators of macrophage fusion. Trends Cell Biol. 19, 514–522 (2009).

57. de Medeiros, V. A. et al. Absence of multinucleated giant cell reaction as an indicator of tumor progression in oral tongue squamous cell carcinoma. Eur. Arch. Oto-Rhino-Laryngol. Off. J. Eur. Fed. Oto-Rhino-Laryngol. Soc. EUFOS Affil. Ger. Soc. Oto-Rhino-Laryngol. - Head Neck Surg. (2021) doi:10.1007/s00405-021-07139-z.

58. Brooks, E., Simmons-Arnold, L., Naud, S., Evans, M. F. & Elhosseiny, A. Multinucleated giant cells’ incidence, immune markers, and significance: a study of 172 cases of papillary thyroid carcinoma. Head Neck Pathol. 3, 95–99 (2009).

59. Lösslein, A. K. et al. Monocyte progenitors give rise to multinucleated giant cells. Nat. Commun. 12, 2027 (2021).

60. Gal-Oz, S. T. et al. ImmGen report: sexual dimorphism in the immune system transcriptome. Nat. Commun. 10, 4295 (2019).

61. Thion, M. S. et al. Microbiome Influences Prenatal and Adult Microglia in a Sex-Specific Manner. Cell 172, 500–516.e16 (2018).

62. Hanamsagar, R. et al. Generation of a microglial developmental index in mice and in humans reveals a sex difference in maturation and immune reactivity. Glia 65, 1504–1520 (2017).

63. Bain, C. C. et al. Rate of replenishment and microenvironment contribute to the sexually dimorphic phenotype and function of peritoneal macrophages. Sci. Immunol. 5, (2020).

64. Klein, S. L. & Flanagan, K. L. Sex differences in immune responses. Nat. Rev. Immunol. 16, 626–638 (2016).

65. Scotland, R. S., Stables, M. J., Madalli, S., Watson, P. & Gilroy, D. W. Sex differences in resident immune cell phenotype underlie more efficient acute inflammatory responses in female mice. Blood 118, 5918–5927 (2011).

66. Mirochnik, Y. et al. Androgen receptor drives cellular senescence. PloS One 7, e31052 (2012).

67. Mirzakhani, K. et al. The androgen receptor-lncRNASAT1-AKT-p15 axis mediates androgen-induced cellular senescence in prostate cancer cells. Oncogene 41, 943–959 (2022).

68. Leng, S. et al. β-Catenin and FGFR2 regulate postnatal rosette-based adrenocortical morphogenesis. Nat. Commun. 11, 1680 (2020).

69. Belenguer, G. et al. RNF43/ZNRF3 loss predisposes to hepatocellular-carcinoma by impairing liver regeneration and altering the liver lipid metabolic ground-state. Nat. Commun. 13, 334 (2022).

70. Bingham, N. C., Verma-Kurvari, S., Parada, L. F. & Parker, K. L. Development of a steroidogenic factor 1/Cre transgenic mouse line. Genesis 44, 419–24 (2006).

71. Xiang, X. et al. TREM2 deficiency reduces the efficacy of immunotherapeutic amyloid clearance. EMBO Mol. Med. 8, 992–1004 (2016).

72. R, S., Ja, F., D, G., Af, S. & A, R. Spatial reconstruction of single-cell gene expression data. Nat. Biotechnol. 33, (2015).

73. Lopez, J. P. et al. Single-cell molecular profiling of all three components of the HPA axis reveals adrenal ABCB1 as a regulator of stress adaptation. Sci. Adv. 7, (2021).

74. Park, S.-Y. & Kim, I.-S. Engulfment signals and the phagocytic machinery for apoptotic cell clearance. Exp. Mol. Med. 49, e331 (2017).

75. Lecoultre, M., Dutoit, V. & Walker, P. R. Phagocytic function of tumor-associated macrophages as a key determinant of tumor progression control: a review. J. Immunother. Cancer 8, e001408 (2020).

76. Janda, E., Boi, L. & Carta, A. R. Microglial Phagocytosis and Its Regulation: A Therapeutic Target in Parkinson’s Disease? Front. Mol. Neurosci. 11, (2018).

77. Kuilman, T. et al. Oncogene-induced senescence relayed by an interleukin-dependent inflammatory network. Cell 133, 1019–1031 (2008).

78. Özcan, S. et al. Unbiased analysis of senescence associated secretory phenotype (SASP) to identify common components following different genotoxic stresses. Aging 8, 1316–1329 (2016).

79. Acosta, J. C. et al. A complex secretory program orchestrated by the inflammasome controls paracrine senescence. Nat. Cell Biol. 15, 978–990 (2013).

80. Fridman, A. L. & Tainsky, M. A. Critical pathways in cellular senescence and immortalization revealed by gene expression profiling. Oncogene 27, 5975–5987 (2008).

81. Coppé, J.-P. et al. Senescence-associated secretory phenotypes reveal cell-nonautonomous functions of oncogenic RAS and the p53 tumor suppressor. PLoS Biol. 6, 2853–2868 (2008).

82. Coppé, J.-P., Desprez, P.-Y., Krtolica, A. & Campisi, J. The senescence-associated secretory phenotype: the dark side of tumor suppression. Annu. Rev. Pathol. 5, 99–118 (2010).

83. Buhl, J. L. et al. The Senescence-associated Secretory Phenotype Mediates Oncogene-induced Senescence in Pediatric Pilocytic Astrocytoma. Clin. Cancer Res. Off. J. Am. Assoc. Cancer Res. 25, 1851–1866 (2019).

84. Saul, D. et al. A New Gene Set Identifies Senescent Cells and Predicts Senescence-Associated Pathways Across Tissues. 2021.12.10.472095 (2021) doi:10.1101/2021.12.10.472095.

85. Newman, A. M. et al. Robust enumeration of cell subsets from tissue expression profiles. Nat. Methods 12, 453–457 (2015).

86. Chen, Z. et al. seq-ImmuCC: Cell-Centric View of Tissue Transcriptome Measuring Cellular Compositions of Immune Microenvironment From Mouse RNA-Seq Data. Front. Immunol. 9, 1286 (2018).

87. Petitprez, F. et al. The murine Microenvironment Cell Population counter method to estimate abundance of tissue-infiltrating immune and stromal cell populations in murine samples using gene expression. Genome Med. 12, 86 (2020).

