## Supplemental Figures 1 to 8 for "Sexually dimorphic activation of innate antitumour immunity prevents adrenocortical carcinoma development"

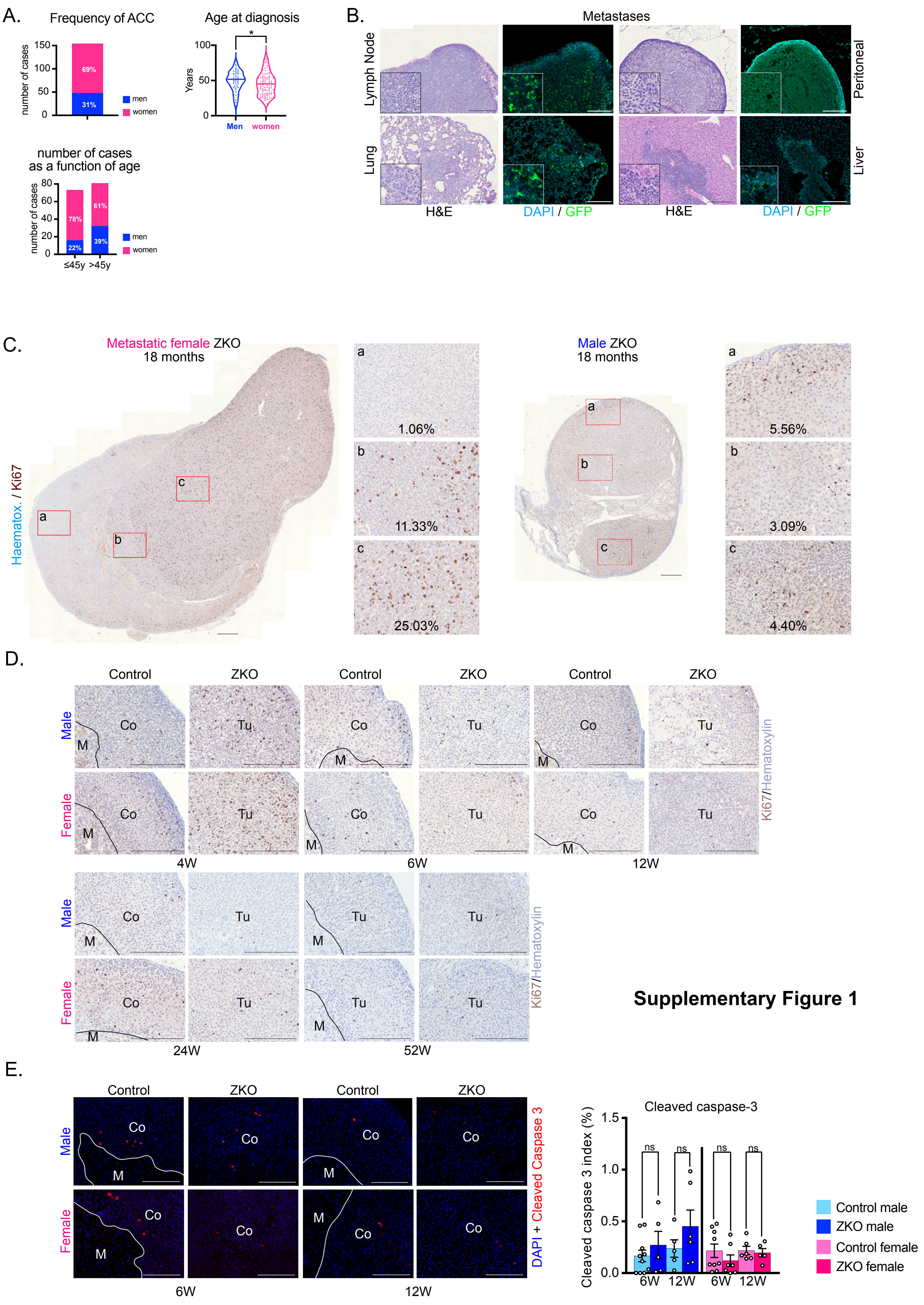


**Supplementary Figure 1**

**Supplementary Figure 1. A-** Demographics of the TCGA cohort of ACC showing overall frequency of ACC in men and women, age at diagnosis as a function of sex and number of cases as a function of age and sex. **B-** Histological (H&E) and immunohistochemical (GFP) analysis of metastases from primary adrenal tumours to lymph nodes, lungs, peritoneal cavity and liver. GFP staining allows identification of tumour cells through SF-1:Cre-mediated recombination of the mTmG locus within the primary tumour. **C-** Immunohistochemical analysis of Ki67 expression in the primary adrenal tumour of 18-month-old *Znrf3 cKO* metastatic female (left) and non-metastatic male (right). The two tumours are represented at the same scale. Insets show heterogeneous proliferation in female tumour and more homogeneous proliferation in male tumour. **D-** Kinetic immunohistochemical analysis of Ki67 expression in control males/females and *Znrf3 cKO* males/females from 4 to 52 weeks. **E-** Immunohistochemical analysis (left panels) and quantification of cleaved-caspase 3 expression (graph) in control males/females and *Znrf3 cKO* males/females at 4 and 12 weeks. Co: cortex; Tu: tumour. Scale bar = 200 µm (B-D); 200µm (E). Graphs represent mean +/- SEM. Statistical analyses in A and E were conducted by Mann-Whitney tests. ns: not significant; * $p<0.05$.

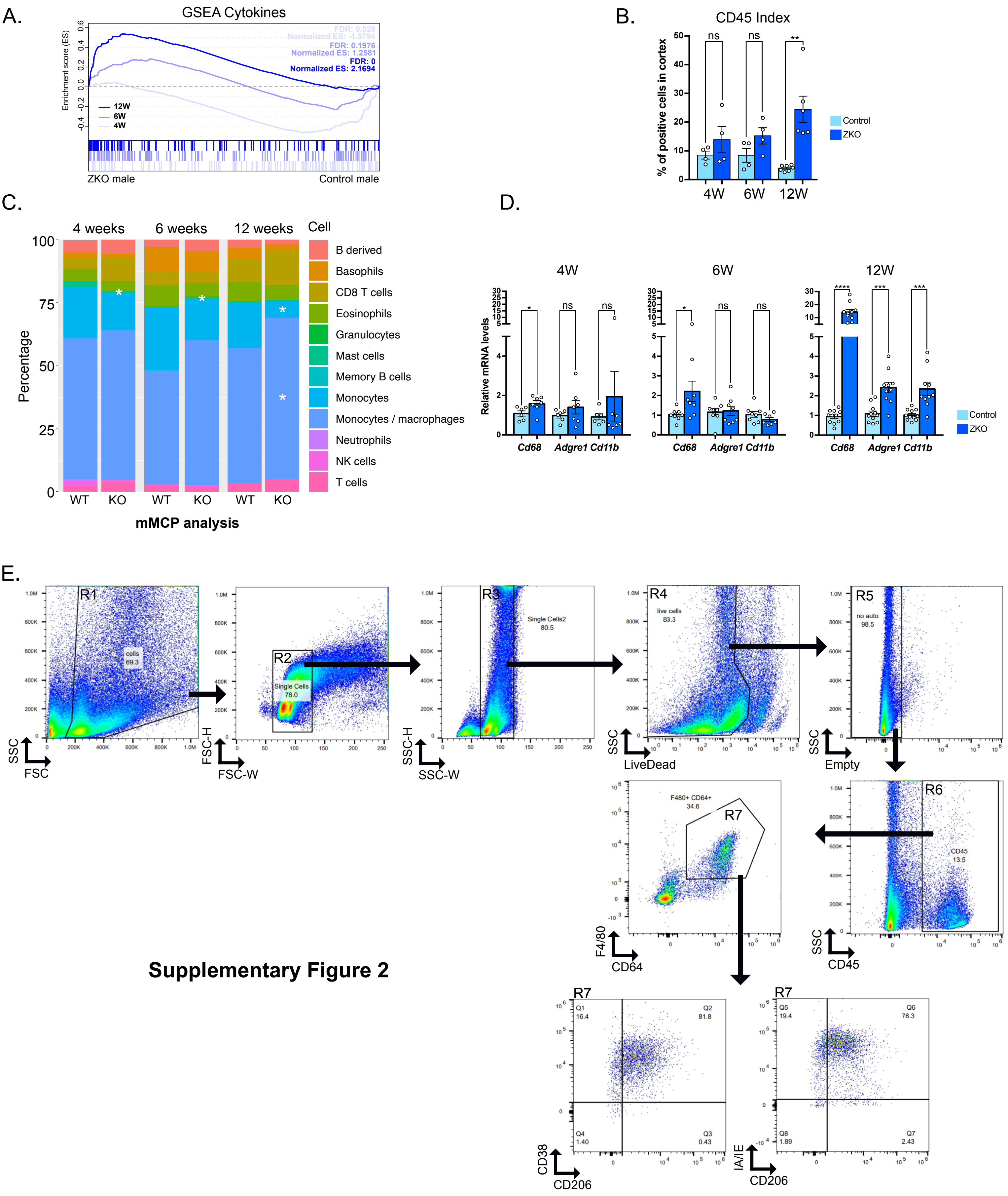


**Supplementary Figure 2**

**Supplementary Figure 2. A-** GSEA of gene expression from 4, 6 and 12 control and *Znrf3 cKO* males. The plot represents enrichment of cytokines gene set in *Znrf3 cKO* compared with controls adrenals. **B-** Quantification of the CD45+ index as the ratio of CD45-positive cells over total nuclei in the cortex of male control and *Znrf3 cKO* mice at 4, 6 and 12 weeks. **C-** Stacked bar plots representing immune cell populations deconvoluted using the mMCP algorithm from gene expression data in control and *Znrf3 cKO* adrenals at 4, 6 and 12 weeks. **D-** RTqPCR analysis of the expression of macrophages-related genes in control and *Znrf3 cKO* males at 4, 6 and 12 weeks. **E-** Flow cytometry gating strategy to identify macrophages in adrenals from control and *Znrf3 cKO* mice. The first gate (R1) was drawn to identify cells of interest and exclude debris. Cells were then gated to exclude doublets (R2 & R3) and live cells were identified using a Live/Dead stain (R4). Live cells were plotted against an empty channel to gate-off auto-fluorescent cells (R5). F4/80+ CD64+ macrophages (R7) were then identified from CD45+ cells (R6). From the macrophage population, M1 macrophage markers CD38 & MHC II (IA/IE) were used against the M2 marker CD206 to identify potential M1 vs M2 polarization. The provided example is from a control male at 4 weeks of age. Graphs represent mean +/- SEM. Statistical analyses in B, C and E were conducted by Mann-Whitney tests. ns: not significant; * $p<0.05$; ** $p<0.01$; *** $p<0.001$; **** $p<0.0001$.

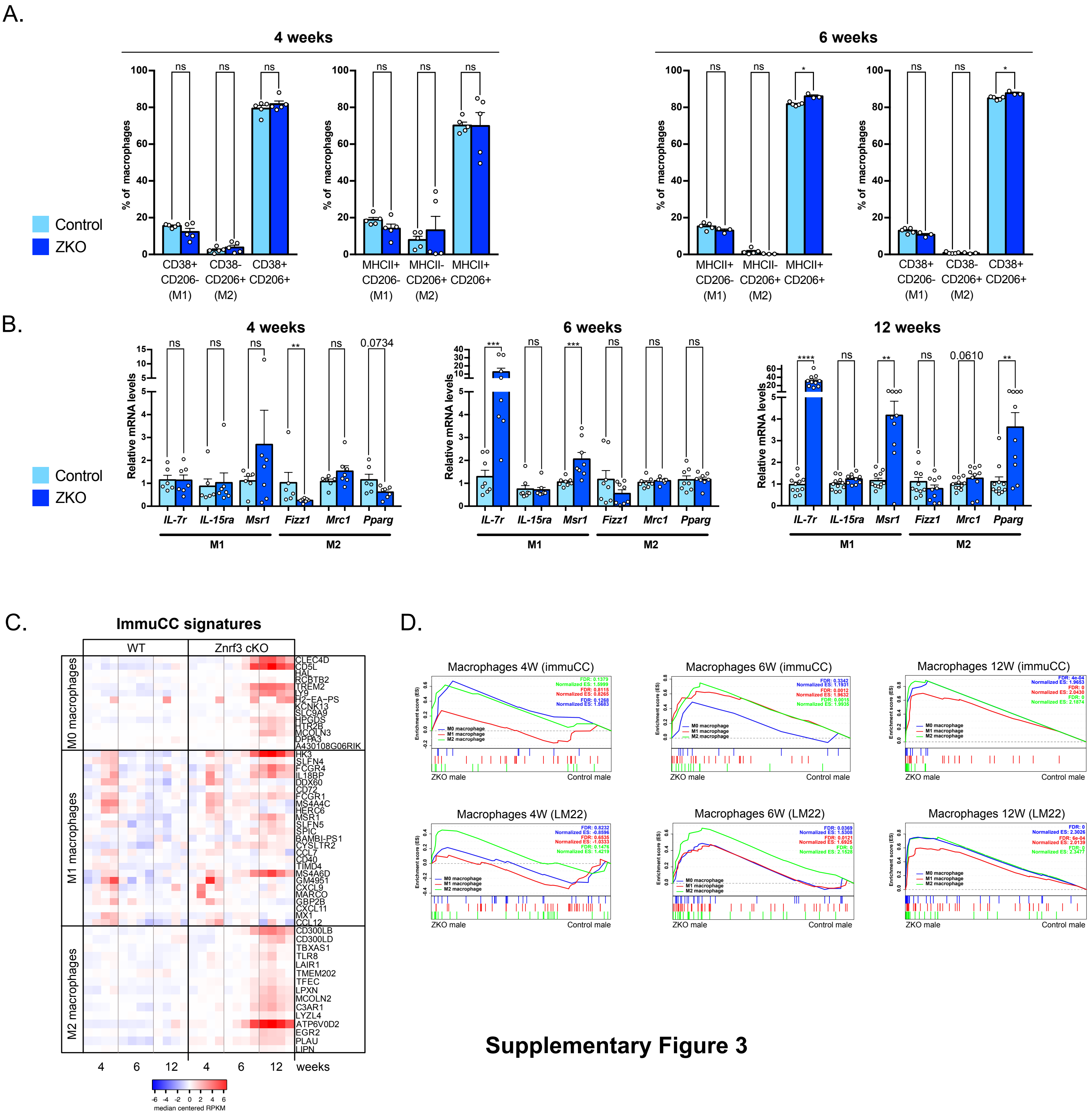


**Supplementary Figure 3**

**Supplementary Figure 3. A-** Quantification of macrophages sub-populations by flow cytometry analysis of control and *Znrf3 cKO* male adrenals at 4 (left panels) and 6 weeks (right panels). **B-** RTqPCR analysis of the expression of M1 and M2 macrophages-related genes in control and *Znrf3 cKO* males at 4, 6 and 12 weeks. **C-** Heatmap showing expression of M0, M1 and M2 gene signatures (extracted from the ImmuCC dataset) in RNA sequencing data from control and *Znrf3 cKO* males at 4, 6 and 12 weeks. All genes in the datasets are represented **D-** GSEA of M0, M1 and M2 macrophages gene sets from the ImmuCC and LM22 datasets in male *Znrf3 cKO* adrenals compared with control adrenals at 4, 6 and 12 weeks. Graphs represent mean +/- SEM. Statistical analyses in A and B were conducted by Mann-Whitney tests. ns: not significant; * $p<0.05$; ** $p<0.01$; *** $p<0.001$; **** $p<0.0001$.

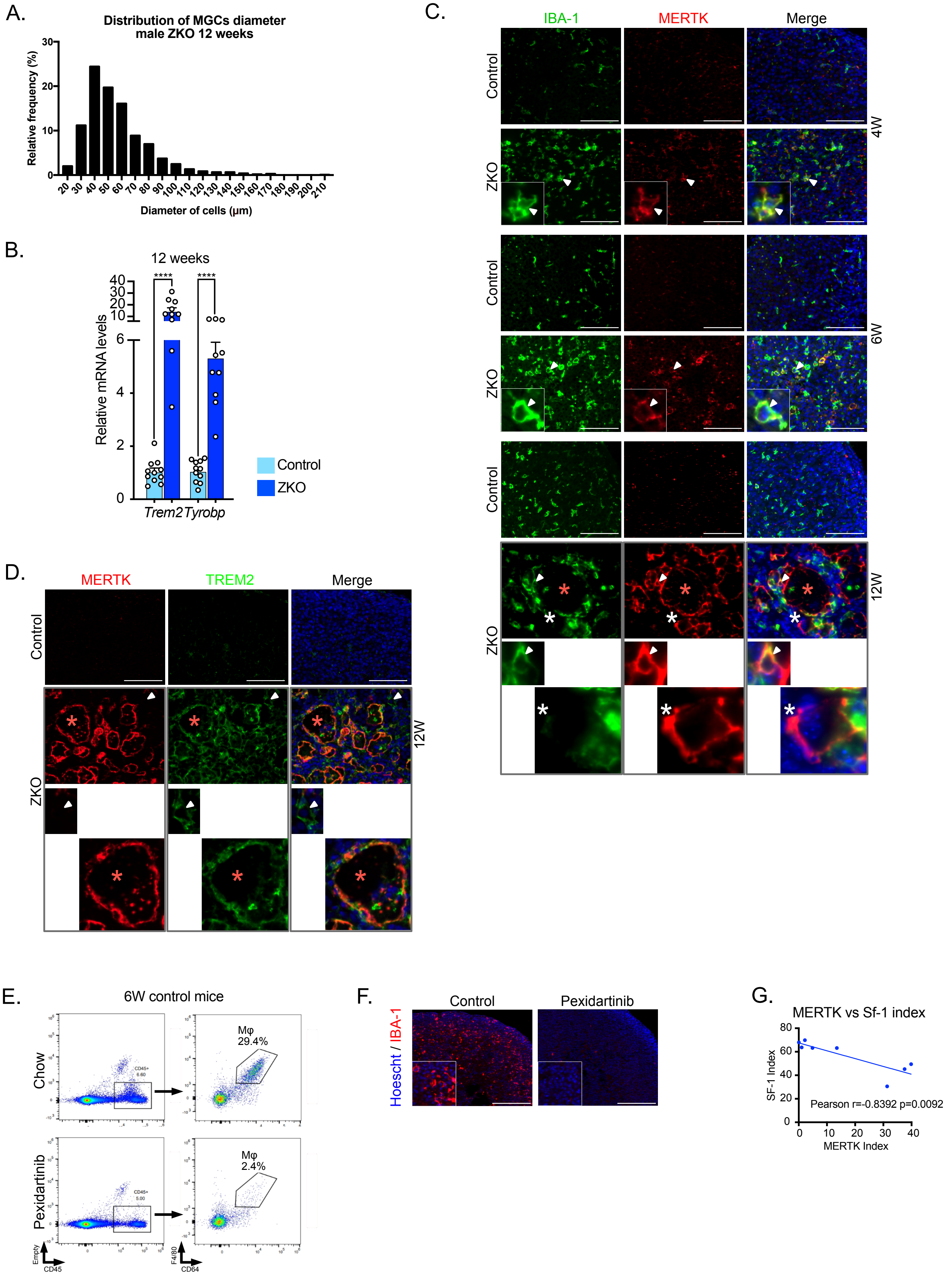


**Supplementary Figure 4**

**Supplementary Figure 4. A-** Diameter distribution of multinucleated giant cells in 12-week-old male *Znrf3 cKO* adrenals (n=5). **B-** RTqPCR analysis of Trem2 and Tyrobp expression in 12-week-old control and *Znrf3 cKO* male adrenals. **C-** Immunohistochemical analysis of IBA-1 and MERTK expression in control and *Znrf3 cKO* adrenals at 4, 6 and 12 weeks. Arrowheads show double-positive macrophages. White stars show MERTK-positive, IBA-1-negative mononucleated macrophages. Red stars show MERTK-positive, IBA-1-negative multinucleated macrophages. **D-** Immunohistochemical analysis of MERTK and TREM2 expression in control and *Znrf3 cKO* adrenals at 12 weeks. Arrowheads show double-positive mononucleated macrophages. Red stars show MERTK-positive, TREM2-positive multinucleated macrophages. **E-** Flow cytometry analysis of CD45+/CD64+/F4/80+ macrophages in the adrenals of 6-week-old control male mice fed for one week with a standard chow (top panels) or a chow enriched with 290 mg/kg of Pexidartinib. **F-** Immunohistochemical analysis of IBA-1 expression in male *Znrf3 cKO* mice treated with either control chow or pexidartinib-enriched chow (290 mg/kg) from 3 to 12 weeks. **G-** Analysis of the correlation between MERTK and SF1-positive cell- indexes in *Znrf3 cKO* mice treated with either control chow or pexidartinib-enriched chow (290 mg/kg) from 3 to 12 weeks. Scale bar = 100 µm (C-D); 200µm (F). Graphs in B represent mean +/- SEM. Statistical analyses in B were conducted by Mann-Whitney tests. **** $p<0.0001$.

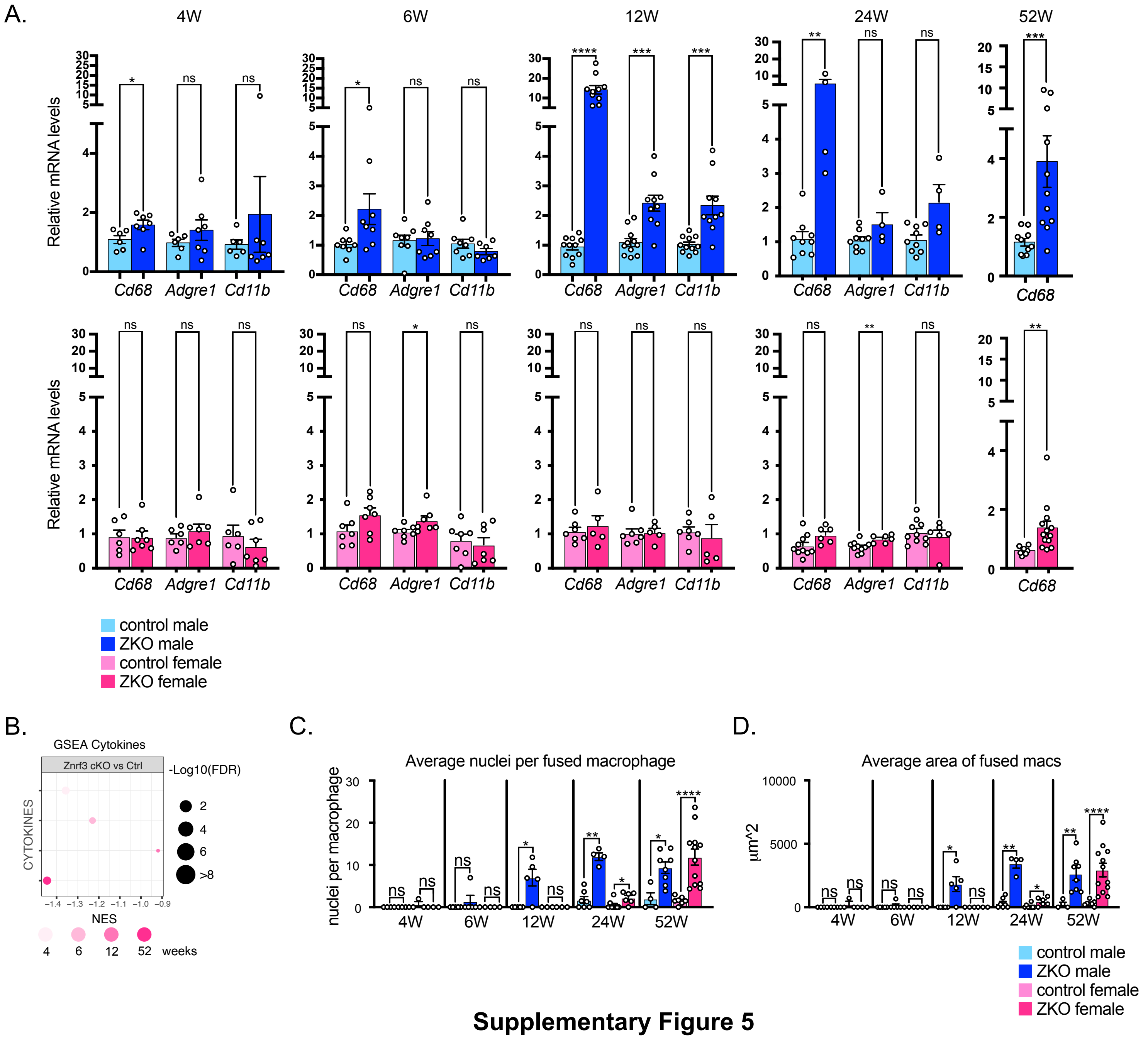


**Supplementary Figure 5**

**Supplementary Figure 5. A-** RTqPCR analysis of the expression of macrophages-associated genes in control and *Znrf3 cKO* males (top panels) and control and *Znrf3 cKO* females (bottom panels) from 4 to 52 weeks. **B-** GSEA of gene expression from 4, 6, 12 and 52-week-old control and *Znrf3 cKO* females. The plot represents enrichment of cytokines gene set in *Znrf3 cKO* compared with control adrenals. **C-** Analysis of the number of nuclei per fused macrophages, determined on H&E sections of control and *Znrf3 cKO* male and female adrenals from 4 to 52 weeks. **D-** Average area of fused macrophages ($\mu m^2$) determined on H&E sections of control and *Znrf3 cKO* male and female adrenals from 4 to 52 weeks. Graphs in A, C and D represent mean +/- SEM. Statistical analyses in A, C and D were conducted by Mann-Whitney tests. ns: not significant; * $p<0.05$; ** $p<0.01$; *** $p<0.001$; **** $p<0.0001$.

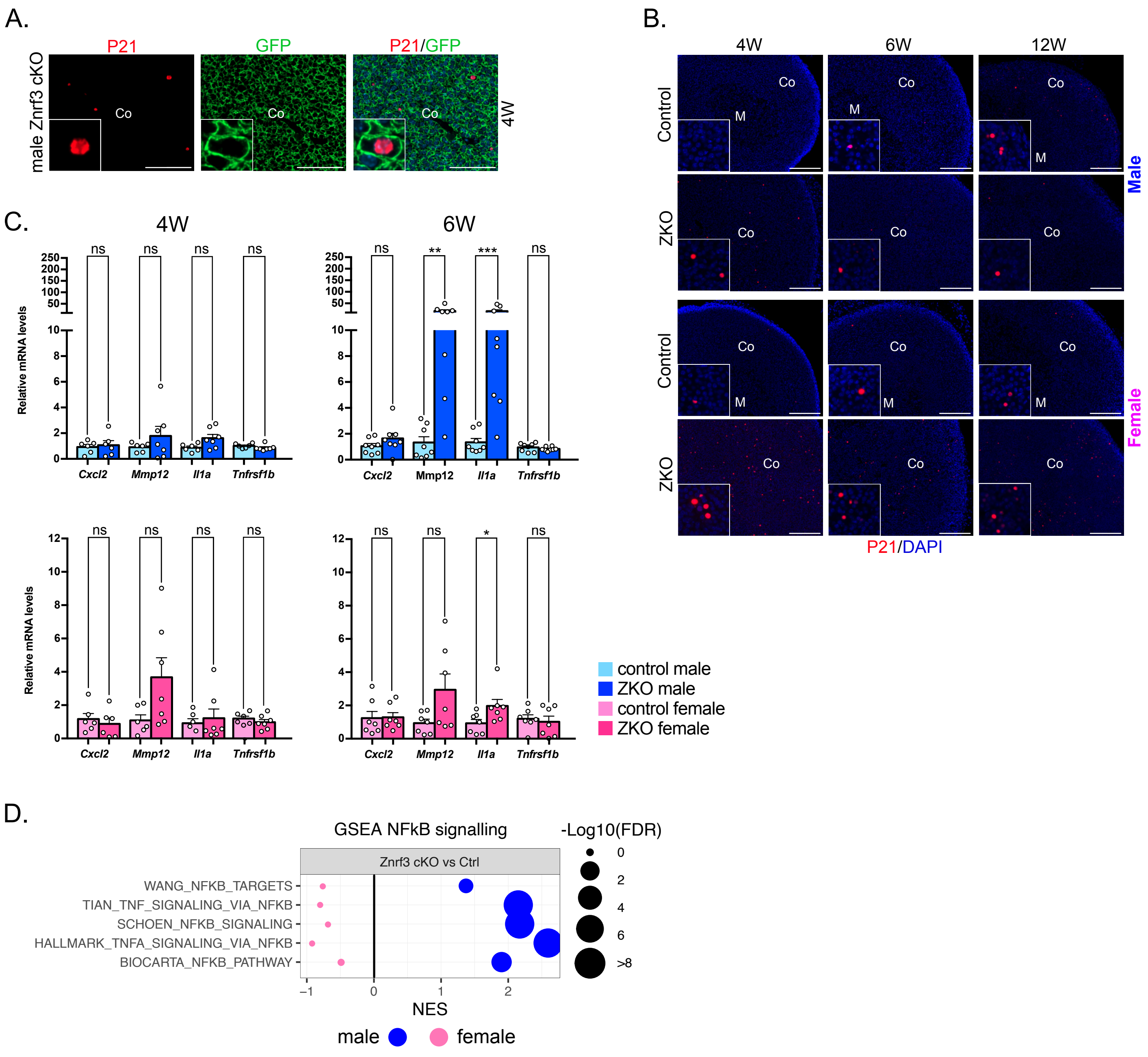


**Supplementary Figure 6**

**Supplementary Figure 6. A-** Immunohistochemical analysis of P21 and GFP (marking SF-1:Cre-mediated recombination of mTmG in steroidogenic cells) in 6-week-old male *Znrf3 cKO* adrenals. **B-** Immunohistochemical analysis of P21 expression in control and *Znrf3 cKO* male and female adrenals at 4, 6 and 12 weeks **C-** RTqPCR analysis of the expression of SASP-associated genes in control and *Znrf3 cKO* males (top panels) and control and *Znrf3 cKO* females (bottom panels) at 4 and 6 weeks. **D-** GSEA of gene expression from 12-week-old control and *Znrf3 cKO* males and females. The plot represents enrichment of NFkB-related gene sets in *Znrf3 cKO* compared with control adrenals (sex matched). Co: cortex; M: medulla. Scale bar = 100 µm (A); 200 µm (B). Graphs in C represent mean +/- SEM. Statistical analyses were conducted by Mann-Whitney tests. ns: not significant; * $p<0.05$; ** $p<0.01$; *** $p<0.001$.

A.

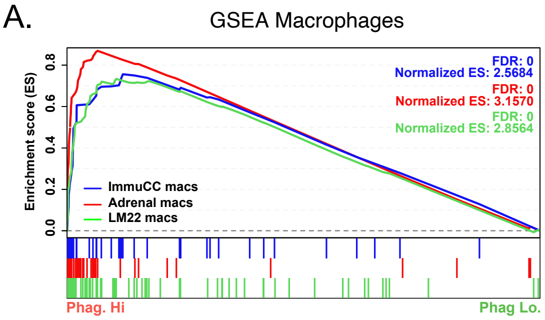


B.

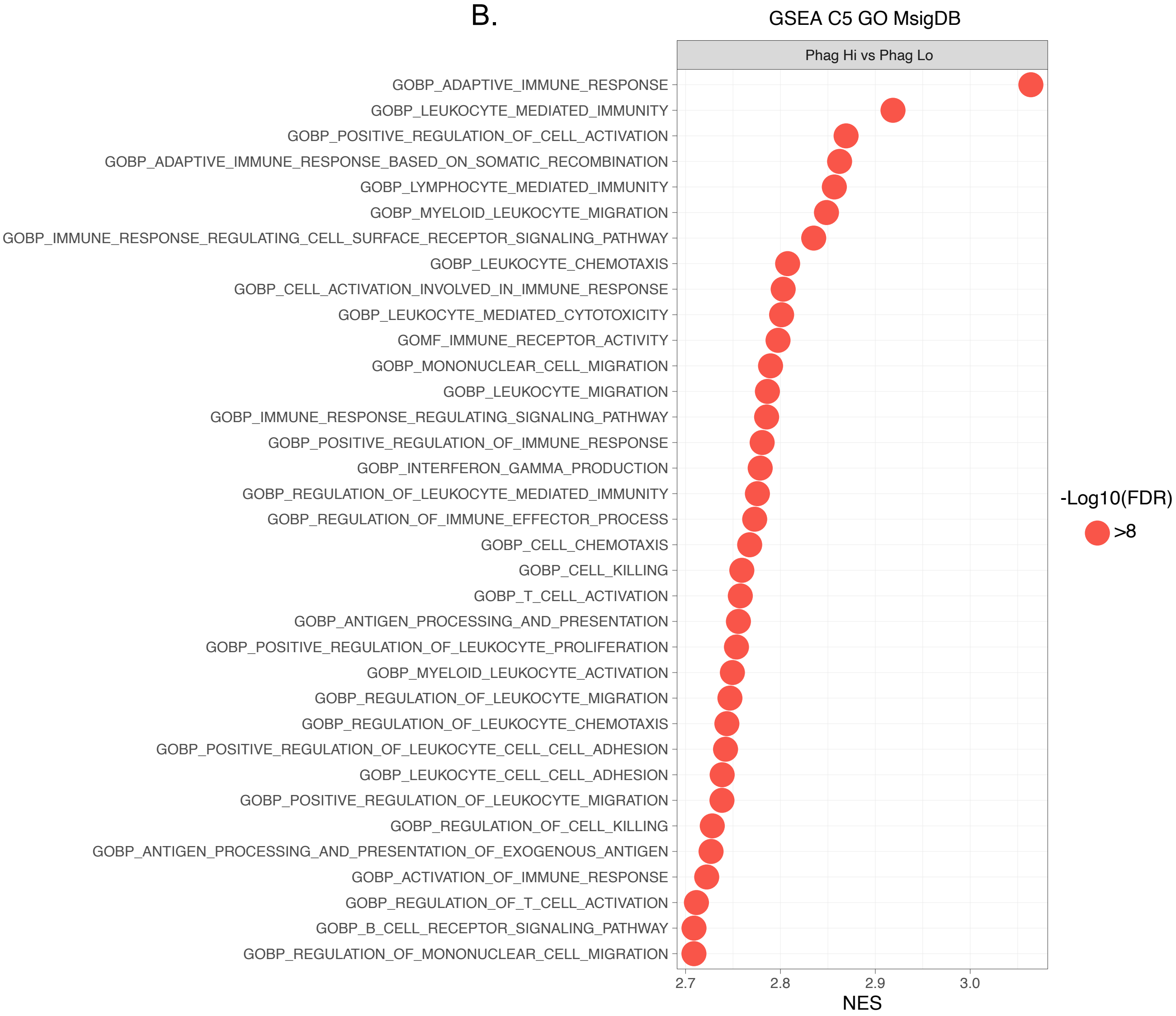


**Supplementary Figure 7**

**Supplementary Figure 7**. **A-** GSEA of macrophages gene sets in TCGA ACC patients with high expression of the phagocytic signature, compared with patients with low phagocytic signature. **B-** GSEA of gene expression from TCGA ACC patients. The plot represents the top 35 enriched gene sets from the C5 Gene Ontology database (MSigDB), in patients with high expression of the phagocytic signature, compared with patients with low phagocytic signature.

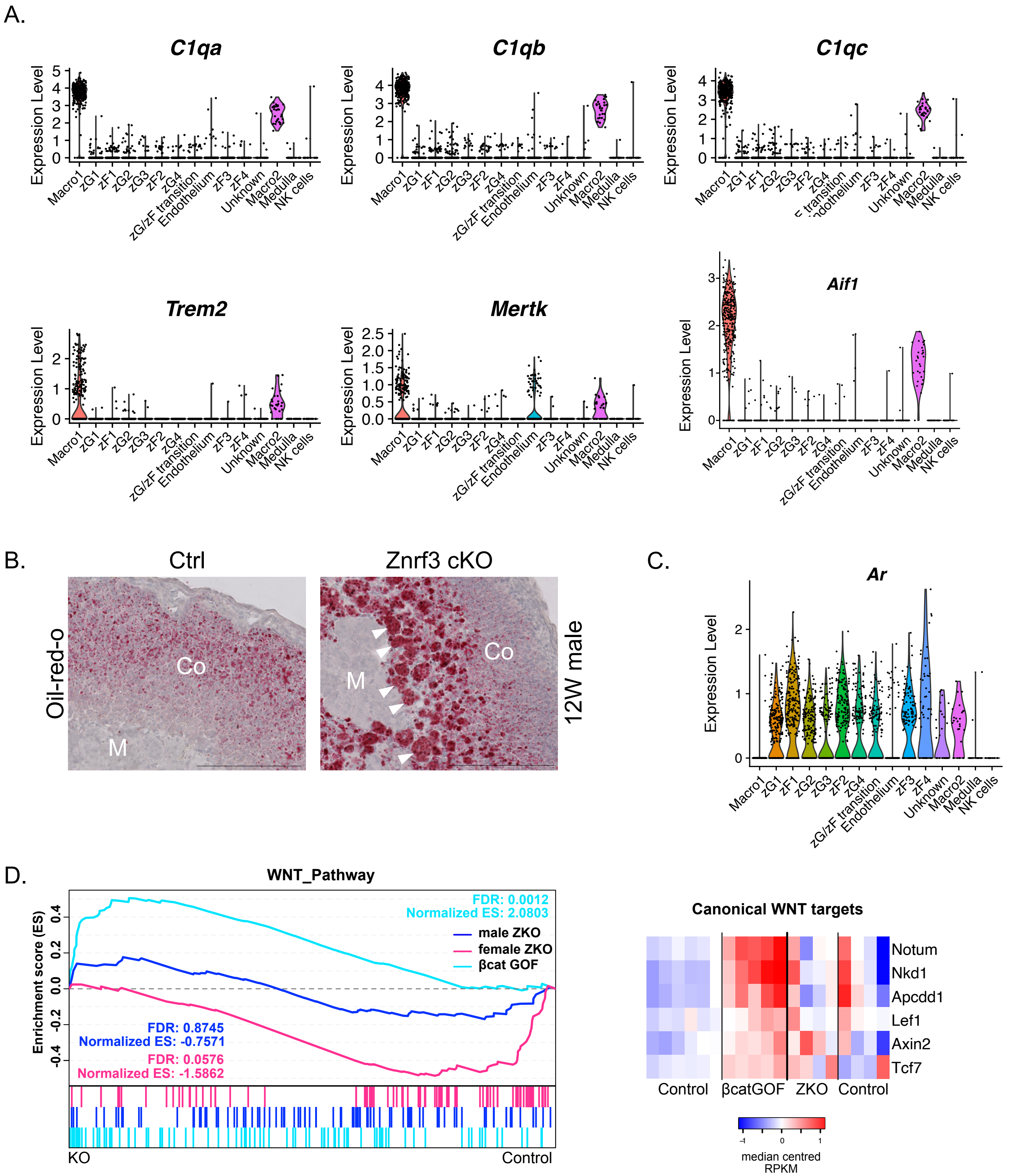


**Supplementary Figure 8**

**Supplementary Figure 8. A-** Expression of macrophages markers in single-cell RNA sequencing data from 10-week-old adult male mouse adrenals **B-** Oil-red-o-staining of 12-week-old control and *Znrf3 cKO* male mouse adrenals. **C-** Expression of *Ar* (androgen receptor*)* in single-cell RNA sequencing data from 10-week-old adult male mouse adrenals **D-** Left panel, GSEA of canonical WNT pathway genes[70] in β-catenin gain of function mice (βcatGOF)[68] and 12-week-old male and female *Znrf3 cKO* mice compared with controls. Right panel, heatmap showing expression of canonical WNT pathway target genes[68] in control, βcatGOF and 12-week-old *Znrf3 cKO* male mice. Arrowheads in B show some of the fused macrophages displaying high levels of lipid accumulation. M: medulla; Co: cortex: Scale bar = 200 μm.
